# Natural gradient enables fast sampling in spiking neural networks

**DOI:** 10.1101/2022.06.03.494680

**Authors:** Paul Masset, Jacob A. Zavatone-Veth, J. Patrick Connor, Venkatesh N. Murthy, Cengiz Pehlevan

## Abstract

For animals to navigate an uncertain world, their brains need to estimate uncertainty at the timescales of sensations and actions. Sampling-based algorithms afford a theoretically-grounded framework for probabilistic inference in neural circuits, but it remains unknown how one can implement fast sampling algorithms in biologically-plausible spiking networks. Here, we propose to leverage the population geometry, controlled by the neural code and the neural dynamics, to implement fast samplers in spiking neural networks. We first show that two classes of spiking samplers—efficient balanced spiking networks that simulate Langevin sampling, and networks with probabilistic spike rules that implement Metropolis-Hastings sampling—can be unified within a common framework. We then show that careful choice of population geometry, corresponding to the natural space of parameters, enables rapid inference of parameters drawn from strongly-correlated high-dimensional distributions in both networks. Our results suggest design principles for algorithms for sampling-based probabilistic inference in spiking neural networks, yielding potential inspiration for neuromorphic computing and testable predictions for neurobiology.

## 1 Introduction

Neural circuits perform probabilistic computations at the sensory, motor and cognitive levels [1–4]. From abstract representations of decision confidence [5] to estimates of sensory uncertainty in visual cortex [6, 7], evidence of probabilistic representations can be found at all levels of the cortical processing hierarchy [8]. To be behaviorally useful, these probabilistic computations must occur at the speed of perception [9]. However, how neuronal dynamics allow brain circuits to represent uncertainty in high-dimensional spaces at perceptual timescales remains unknown [3, 10–12].

Several neural architectures for probabilistic computation have been proposed, including: probabilistic population codes [13], which in certain cases allow a direct readout of uncertainty; direct encoding of metacognitive variables, such as decision confidence [4, 5, 8]; doubly distributional codes [14, 15] which distinguish uncertainty from multiplicity; and sampling-based codes [9, 16–25], where the variability in neural dynamics corresponds to a signature of exploration of the posterior probability. Most experiments quantifying uncertainty representations in single biological neurons have only varied parameters along one or two dimensions, such as in Bayesian cue combination [2, 10, 26]. In these conditions, many algorithms can perform adequately. However, probabilistic inference becomes more challenging as the entropy of the posterior distribution—which often scales with dimensionality—increases [27, 28]. Some algorithms that work well in low dimensions, such as probabilistic population codes, may scale poorly to high-dimensional settings [3].

Of the proposed approaches to probabilistic computation in neural networks, sampling-based codes are grounded in the strongest theoretical framework [27, 29–33], and have been used to perform inference at scale [34–36]. Moreover, they predict specific properties of neural responses in visual cortex, including changes in Fano factor and frequency of oscillations with tuning and stimulus intensity [23]. Previous works have proposed several approaches to accelerate sampling in biologically-inspired algorithms. Hennequin et al. [9] showed that adding non-reversible dynamics to rate networks can reduce the sample autocorrelation time. However, they did not study convergence of the sampling distribution, and did not consider the biologically-relevant setting of spiking networks. Savin and Denève [20] used a distributed code to parallelize sampling in spiking networks, but only considered two-dimensional distributions. Therefore, it remains unclear how accurate sampling from high-dimensional distributions at behaviorally-relevant timescales can be achieved using spiking networks.

In this paper, we show how the choice of the geometry of neural representations at the population level [37, 38], set by the neural code and the neural dynamics, can accelerate sampling-based inference in spiking neural networks. Ideas from information geometry allow us to perform inference in the natural space of parameters, which is a manifold with distances measured by the Fisher-Rao metric (Figure 1.c) [39–42]. Concretely, we leverage recently-proposed methods for accelerating sampling from the machine learning literature [28, 42–44] to design novel efficient samplers in spiking neural networks. The structure and major contributions of this paper are divided as follows:

- In §2, we construct from first principles a novel spiking neural network model for sampling from multivariate Gaussian distributions. This model is based on a probabilistic spike rule that implements approximate Metropolis-Hastings sampling. We show that efficient balanced networks (EBNs) [20, 45–49] emerges as a limit of this model in which spiking becomes deterministic.
- In §3, we show that population geometry enables rapid sampling in spiking networks. Leveraging the “complete recipe” for stochastic gradient MCMC [43], we establish principles for the design of efficient samplers in spiking neural networks. Then, we show how neural population geometry enables fast sampling—on the timescale of tens of milliseconds—in two limits of the model introduced in §2: EBNs in which sampling is driven by stochastic Langevin dynamics [20], and networks in which sampling is driven purely by a Metropolis-Hastings probabilistic spiking rule.
- Finally, in §4 we conclude by discussing the implications of our results in the context of prior works, and highlight their limitations as well as remaining open questions. In particular, we comment on possibilities for future experimental studies of sampling in biological spiking networks, and applications to neuromorphic computing.

**Figure 1:**
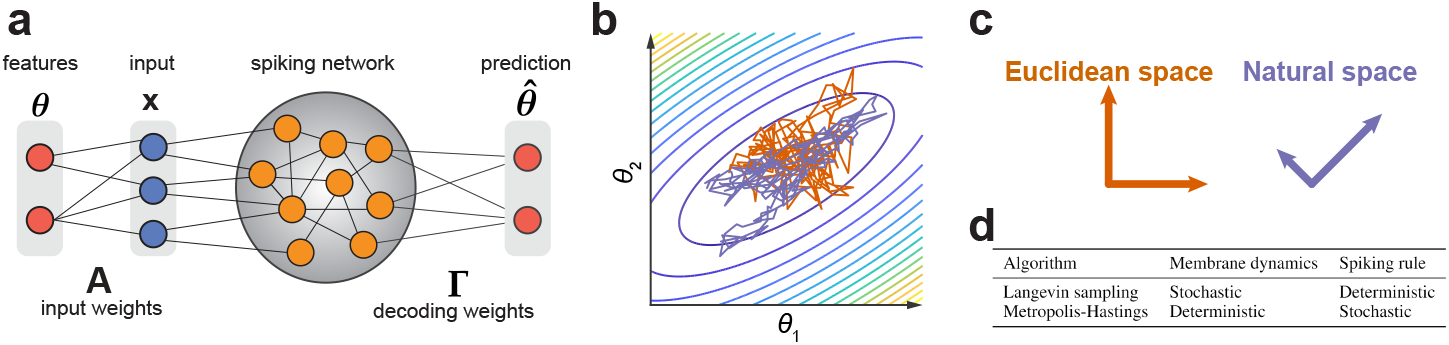
Probabilistic inference in spiking neural networks.**a**. Circuit diagram. **b**. Langevin sampling in Euclidean space and natural space. **c**. Geometry of the inference space in (b). Arrows indicate the principal directions of the sampling noise covariance. **d**. Comparison of Langevin and Metropolis-Hastings sampling algorithms.

## 2 Spiking networks for sampling-based probabilistic inference

We begin by proposing a framework for probabilistic inference in spiking neural networks in which the spiking rule implements a Metropolis-Hastings step. We show that EBNs [20, 45–49] can be recovered as a limiting case of this more general framework.

In this work, we will keep our discussion quite general, and state our results for sampling from a generic Gaussian distribution. However, the problem we aim to solve can be given a concrete interpretation in a neuroscience context, and could also be extended to non-Gaussian distributions. The goal of a neural network performing probabilistic inference is to estimate a posterior distribution *P* (***θ***| **x**) over *n*_*p*_ latent variables (or parameters) ***θ*** given an input **x** (Figure 1). The input could correspond to the activity of sensory neurons in early sensory processing (e.g. input onto ganglion cells in the retina or onto mitral cells in the olfactory bulb) or inputs into a cortical column that linearly sense features in the environment through an affinity matrix. We provide a detailed discussion of this linear Gaussian model in Appendix C.2. In the rest of the paper, we will usually abbreviate the distribution from which we want to sample as *P* (***θ***), rather than writing *P* (***θ***|**x**).

### 2.1 Deriving an approximate Metropolis-Hasting spiking sampler

We will build a network of *n*_*n*_ spiking neurons to approximately sample an *n*_*p*_-dimensional Gaussian distribution of time-varying mean ***θ***(*t*) and fixed covariance **Ψ**. We first consider the case in which ***θ*** is constant, and then generalize the resulting algorithm to the case in which it is slowly time-varying.

As in prior work on probabilistic inference using spiking networks, we take the samples **z** to be linearly decoded from the filtered spike trains **r** of *n*_*n*_ neurons [20, 45–49]. Working in discrete time for convenience and clarity (as in [48]), we let

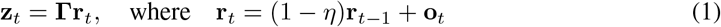

**z**_*t*_ = **Γr**_*t*_, where **r**_*t*_ = (1 − *η*)**r**_*t*−1_ + **o**_*t*_ (1) is the low-pass filtered history of spikes 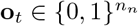 for some decay constant 0 ≤ *η* ≤ 1.

Metropolis-Hastings sampling constructs a Markov chain by drawing a proposed next state from some distribution, and then deciding whether to accept or reject that proposal based on a probabilistic rule [27, 50]. The acceptance ratio is given in terms of the relative posterior probability of the proposed and current states. Here, we will randomly propose which neuron spikes at a given timestep.

In trying to build a Metropolis-Hastings sampler [27, 50] using a probabilistic spiking rule, we are immediately faced with two problems. First, the spikes are sign-constrained and discrete. Second, the dynamics of the filtered spike history incorporate a decay term, hence the readout **z** will change even if no spikes are emitted. These conditions mean that the proposal density over **z** will not be symmetric, and that the Markov process will not satisfy the condition of detailed balance [27, 50]. We note that previous work has shown that violations of detailed balance can accelerate sampling [17, 18, 51], but we will not carefully explore this possibility in the present work. To obtain a symmetric proposal distribution with sign-constrained spikes, we assume that the network is divided into two equally-sized populations with equal and opposite readout weights, i.e. that the readout matrix is of the form **Γ** = [+**M, M**] for some matrix 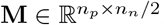. This could be accomplished by dividing the total population of neurons into excitatory and inhibitory populations with weights that are fine-tuned to be equal and opposite. The second problem can be solved by assuming that *η* = 0, i.e., that we have access to a perfect integrator of the spike trains.

At the *t*-th timestep, we choose one neuron *j* uniformly at random, and let the spike proposal be **o**^*′*^ = **e**_*j*_, where **e**_*j*_ is the *j*-th standard Euclidean basis vector (i.e., (**e**_*j*_)_*k*_ = *δ*_*jk*_). This yields a candidate readout

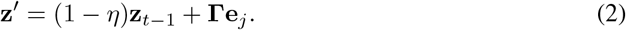

If *η* = 0 and the balance assumption on **Γ** is satisfied, then the proposal distribution is exactly symmetric, in the sense that the probabilities of reaching **z**^*′*^ from **z**_*t*−1_ and of reaching **z**_*t*−1_ from **z**^*′*^ are equal. Then, if we accept the proposed spike with probability

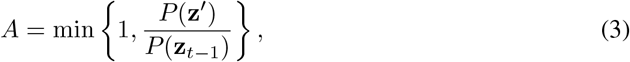

we obtain a Metropolis-Hastings sampling algorithm for the discretized Gaussian [27, 50].

We now relax the assumption of the perfect integration, and assume only that *η* ≪ 1. Introducing decay smooths the readout, which in the perfect integrator is discretized. With the same proposal distribution as before, we take the acceptance ratio of the accept-reject step to be

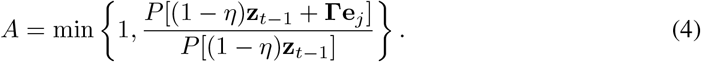

This choice has two important features. First, the decay means that the proposal distribution will be asymmetric, and the Markov chain will no longer satisfy the condition of detailed balance [27, 50]. However, the resulting error will be small if *η* ≪ 1. Second, by comparing the likelihood of the proposal, *P* [(1 −*η*)**z**_*t*− 1_ + **Γe**_*j*_], to the likelihood of the next state without the proposed spike but with the decay *P* [(1− *η*)**z**_*t*− 1_] (instead of the likelihood of the current state *P* [**z**_*t*− 1_] as in the Metropolis-Hastings algorithm), this choice implements a sort of look-ahead step that should allow the algorithm to partially compensate for the decay in the rate.

With the choices above, we show in Appendix B.2 that one can write the acceptance ratio (4) as

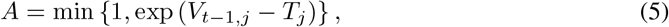

where **V**_*t*−1_ = −(1 − *η*)**Ωr**_*t*−1_ + **Γ**^⊤^**Ψ**^−1^***θ*** has the interpretation of a membrane potential,

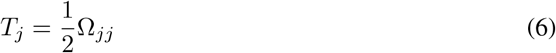

has the interpretation of a spiking threshold, and the the recurrent weight matrix is defined as

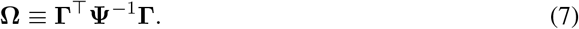

Thus far, we have assumed that the mean signal is constant. The natural generalization of this algorithm to a time-varying mean signal ***θ***_*t*_ is to take the membrane potential to be **V**_*t*_ = − (1 −*η*)**Ωr**_*t*_ + **Γ**^⊤^**Ψ**^−1^***θ***_*t*_. This leads to the voltage dynamics

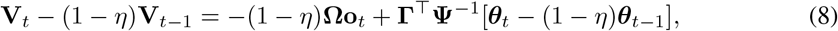

which, when combined with the probabilistic spiking rule with uniform proposals and acceptance ratio (5), yields our final algorithm. This will not be an exact Metropolis-Hastings sampler unless the mean is constant, the decay term vanishes (*η* = 0), and the readout matrix **Γ** satisfies an exact balancing condition. In particular, if these conditions are violated, the Markov chain will not satisfy detailed balance. However, if they are violated only weakly, this algorithm can still be a reasonable approximation to a true sampler. We will provide empirical evidence for this intuition in §3.

### 2.2 The continuous-time limit

We now consider the continuous-time limit of the model introduced above. This limit corresponds to taking the limit in which spike proposals are made infinitely often, and regarding the dynamics written down previously as a forward Euler discretization of an underlying continuous-time system. For a timestep Δ between spike proposals, we let the discrete-time decay rate be *η* = Δ*/𝒯*_*m*_ for a time constant *𝒯*_*m*_, thusly named because it has the interpretation of a membrane time constant. Then, we show in Appendix B.4 that the Δ↓ 0 limit of the discretized rate dynamics (1) yields the familiar continuous-time dynamics

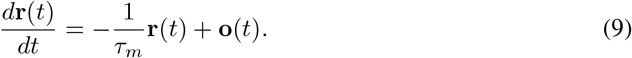

In continuous time, the spike train **o**(*t*) is now composed of Dirac delta functions, as the discretized spikes are rectangular pulses of width Δ in time and height 1*/*Δ. We next consider the voltage dynamics of the leaky integrator for a varying mean signal (8), which have a similar continuum limit:

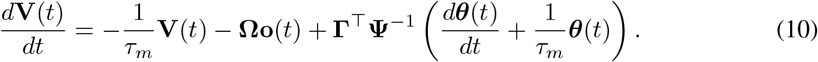

In this limit, the rate will decay by only a infinitesimal amount between a rejected spike proposal and the next proposal, meaning that the error incurred by neglecting the asymmetry in the proposal distribution due to the decay should be negligible. We also note that, though the probabilistic spike rule (5) does not explicitly include a reset step, the dynamics (10) prescribe that the *j*-th neuron’s membrane voltage should be decremented by 2*T*_*j*_ after it spikes.

### 2.3 Efficient balanced networks as a limit of the spiking Metropolis-Hastings sampler

This spiking network samples the posterior distribution by emitting spikes probabilistically, but we can use the same architecture to re-derive EBNs, which approximate continuous dynamical systems using spiking networks [45–49]. If we take 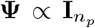 (for 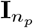 the *n*_*p*_ × *n*_*p*_ identity matrix) the voltage dynamics (10) are identical to those of the EBN^3^. The greedy spiking rule of the EBN can be recovered in this framework by taking a limit in which the variance of the Gaussian target distribution vanishes. Concretely, we let 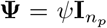, and define re-scaled variables 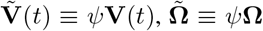, and 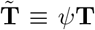 that will remain 𝒪 (1) even as we take *ψ* ↓ 0. In terms of these new variables, the acceptance ratio is 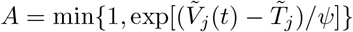, which tends to 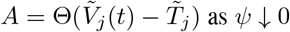 as *ψ*↓ 0. This explicitly recovers the greedy spike rule used in EBNs. In Appendix B, we further analyze how the overall scales of **Ψ** and **Γ** affect the probabilistic spike rule.

The network with voltage dynamics (10) samples a distribution with mean ***θ*** and covariance **Ψ**. Instead of sampling using the structured proposal distribution on spikes, this network can implement sampling through slowly varying Langevin dynamics on ***θ***. In the limit where the spike rule becomes greedy, this recovers the spiking sampler studied by Savin and Denève [20]. We will discuss this model further in §3.

## 3 Population geometry for fast sampling

### 3.1 Leveraging the geometry of inference to accelerate sampling

We first review recent work from the machine learning literature for how population geometry can be chosen to accelerate simple Langevin sampling, which establishes principles for the design of fast samplers. For probability distributions belonging to the exponential family, including the Gaussian distributions on which we focus in this work, one can write the density *P* (**z**) in terms of an energy function *U* (**z**) such that *P* (**z**) ∝ exp[−*U* (**z**)]. The classic algorithm to sample such a distribution is the discretization of the naïve Langevin dynamics

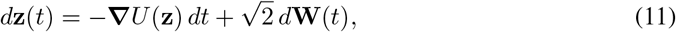

where **W**(*t*) is a standard Brownian motion [29, 30, 33, 52]. By simply following these dynamics, one can obtain samples from the target distribution and therefore an estimate of the uncertainty at the timescale taken by the network to sufficiently explore the target distribution.

The Langevin dynamics (11) can be directly implemented in a rate network [9, 19] or approximately implemented in a spiking network [20], but their convergence properties for high-dimensional distributions have not been investigated in a neuroscience context. It is well known in statistics that, as the dimensionality of the target distribution increases, convergence of Langevin sampling to the target distribution slows dramatically. Furthermore, the discretization step can induce errors that cause the variance estimated from sampling to exceed the target variance [29, 30, 33, 52].

To overcome these issues, prior work in statistics and machine learning has proposed algorithms that can accelerate the sampling [28, 31, 42–44, 53–55] which we leverage here to propose our fast samplers in spiking neural networks. These ideas were unified into a common framework by Ma et al. [43] (and see also [44, 56]) who proposed a “complete recipe” for stochastic gradient MCMC:

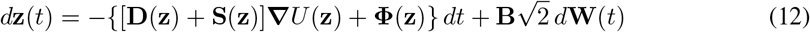

with **BB**^⊤^ = **D**. The matrix fields **D**(**z**) and **S**(**z**) modify the dynamics but keep the target distribution unchanged. **D** is positive semi-definite and defines the local geometry of the space in which the inference is occurring, while **S** is skew-symmetric and adds non-reversible dynamics. When **D** and **S** are state dependent, the correction term **Φ** = div(**D** + **S**) must be included [43].

The “complete recipe” provides a general framework to design samplers based on Langevin dynamics. Samplers based on Riemannian geometry can be designed by choosing **D** to be the inverse of the Fisher information matrix **G** (or an approximation thereof), yielding the natural gradient **Δ**_nat_*U* = **G**^−1^ **Δ***U* (Figure 1) [39–42, 57, 58]. Samplers incorporating dummy variables can be designed by expanding the parameter space and using the matrices **D** and **S** to obtain the desired dynamics [31, 43, 44, 53–55, 59]. Similarly, prior works have proposed methods to accelerate sampling in biologically inspired neural networks by parallelizing the inference [20], using Hamiltonian dynamics [21] or by adding non-reversible dynamics [9, 17, 18]. The “complete recipe” provides a general framework that encompasses all these examples, allowing for the principled design of biologically-plausible samplers.

In the simple setting of linear rate networks sampling from a Gaussian distribution with constant mean ***μ*** and covariance **∑**, the complete recipe for state-independent **D** and **S** takes the form

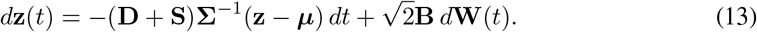

In Appendix D, we show in the zero-mean case ***μ*** = **0** that large eigenvalues of the covariance matrix **∑** introduce long timescales in the sample cross-covariance for naïve Langevin dynamics 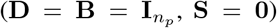, which slow convergence to the target distribution. Concretely, the 2-Wasserstein distance between the ensemble sampling distribution at time *t* and the target is

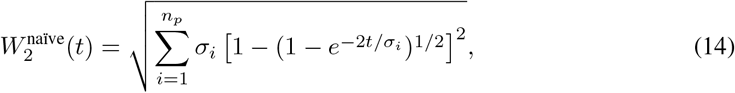

where *σ*_*i*_ are the eigenvalues of the covariance matrix. Networks performing inference in the natural space (**D** = **∑, B** = **∑**^1*/*2^) are insensitive to these large eigenvalues, with 2-Wasserstein distance

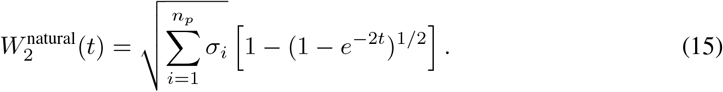

For this analytical result, we consider the distribution of samples at time *t* over realizations of the noise process, which differs from the distribution of samples over time within a single realization, as used elsewhere in the paper. This choice is made because the joint distribution of samples within a single realization is challenging to characterize [60]. These results show how natural gradient can accelerate inference in linear rate networks, and complement Hennequin et al. [9]’s study of how adding irreversible dynamics can reduce the sample autocorrelation time (i.e.,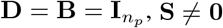).

### 3.2 Fast sampling through population geometry in efficient balanced networks

We first consider a sampler based on efficient neural networks [20, 45–47] that leverages the geometry of the inference to implement efficient sampling at the level of the population. In previous work, Savin and Denève [20] derived a sampler implementing naïve Langevin dynamics (we provide a full derivation using the notation from the present paper in Appendix C). Although they proposed to accelerate sampling by implementing parallel inference loops, they do not leverage the geometry of the inference nor do they test the convergence in high (*n*_*p*_ > 2) dimensions.

Here, we approximate the “complete recipe” dynamics (13) using an EBN, and show that performing inference using natural gradients helps with speed and accuracy in high dimensions. As in §3, we consider sampling from a Gaussian distribution of possibly time-varying mean ***μ*** and covariance **∑**.^4^ As we principally study the effect of the geometry, we henceforth set **S** = **0**. Note that even though the underlying dynamics of (13) are reversible if **S** = **0**, the non-linearities introduced by spiking lead to a non-reversible sampler. To embed these dynamics in the EBN framework, we take the target mean in the voltage dynamics (10) to evolve according to the Ornstein-Uhlenbeck process (13), which we generalize to include an intrinsic timescale *𝒯*_*s*_. Following [20], we approximate the true state of the sampling dynamics by the instantaneous estimate from the filtered spike history. Then, as detailed in Appendix C, we obtain an EBN with the voltage dynamics

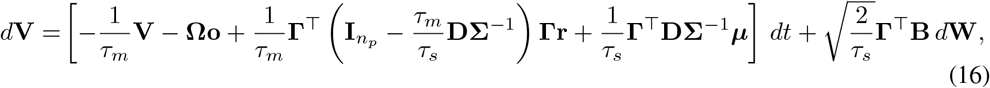

where the recurrent weight matrix is **Ω** = **Γ**^⊤^**Γ**, and the spiking rule is greedy, with thresholds *T*_*j*_ = (**Γ**^⊤^**Γ**)_*jj*_*/*2.

To illustrate how correlations between parameters affect sampling in high dimensions, we will focus on equicorrelated multivariate distributions. The covariance matrix **∑** of such a distribution is parameterized by an overall variance *σ* > 0 and a correlation coefficient −1 < *ρ* < 1 such that ∑_*ij*_ = *σ*[*δ*_*ij*_ + *ρ*(1 −*δ*_*ij*_)]. We will explore the performance of the sampling algorithms across values of *ρ* for different dimensions of the parameter (*n*_*p*_) and neuron (*n*_*n*_) spaces. For a multivariate Gaussian distribution 𝒩 (***μ***,**∑**), the Fisher information matrix is **G** = **∑**^−1^ and we will therefore use **D** = **G**^−1^ = **∑** (and 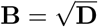) in our geometry aware implementation (see Appendix C.3). In our simulations, we compare the accuracy of sampling of the naïve implementation (**D** = **B** = **I**) with the geometry-aware version over a 50 millisecond window, which is roughly twice the membrane time constant *𝒯*_*m*_ = 20 ms, as well as at steady-state. In Figures 2, E.1 and E.3, we show that the geometry-aware sampler is more robust to increasing the correlation of the parameters and the dimensionality, allowing inference at behavioral timescales. As in non-spiking Langevin sampling, discretization leads to an overestimation of variance [27], but this effect, although still present, is strongly reduced in the geometry-aware implementation. Note here, that the geometry is imposed through the dynamics of the membrane potentials via the recurrent connectivity in the network.

**Figure 2:**
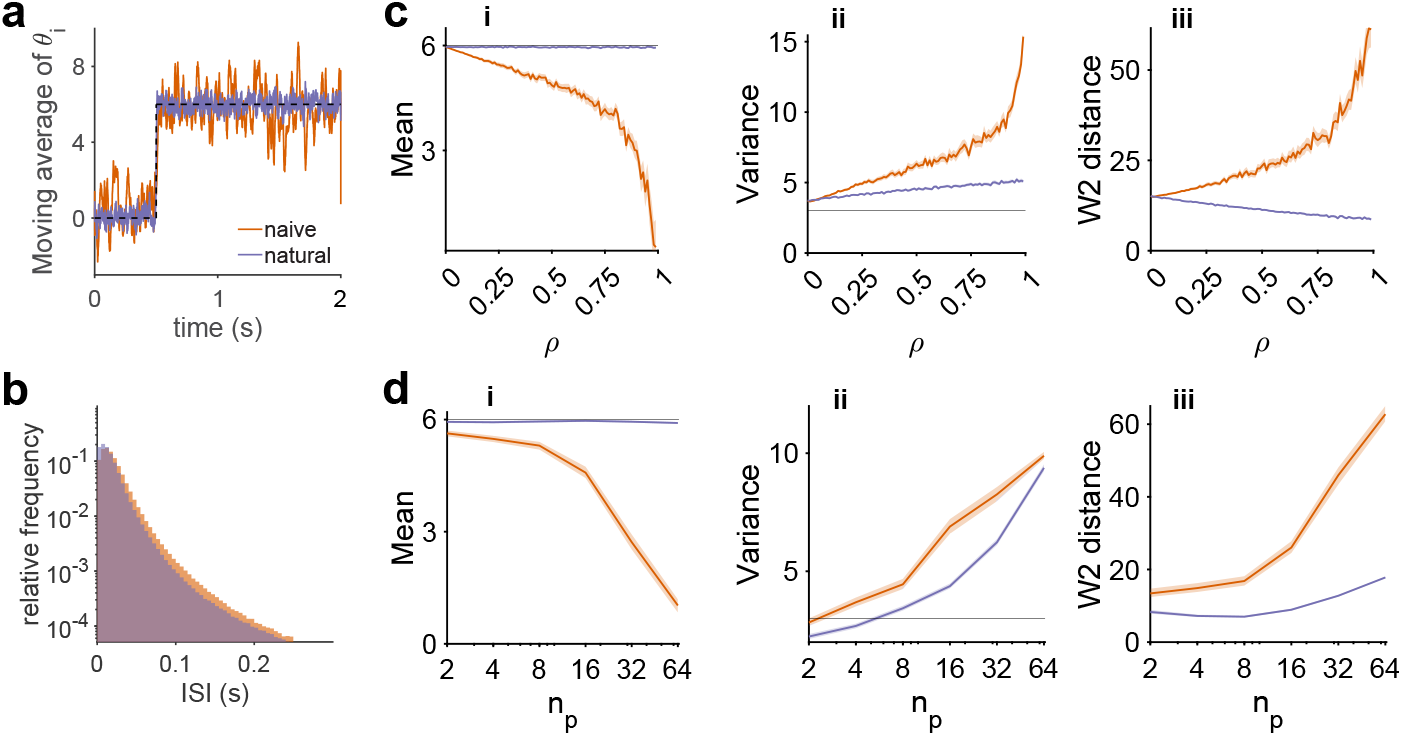
Accelerating inference through population geometry in EBNs simulating Langevin sampling: 50 ms moving averages of parameter estimates 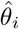 over time, for naïve and natural geometry in a network sampling from *n*_*p*_ = 20-dimensional equicorrelated Gaussian distribution of correlation *ρ* = 0.75 using a network of *n*_*n*_ = 200 neurons. At *t* = 0.5 s, the target mean shifts from zero to six, as indicated by the black dashed line. **b**. The distribution of ISIs across neurons and trials is approximately exponential. See Figure E.2 for more statistics of the resulting spike trains. **c**. Comparison of performance in the 50 ms following stimulus onset (after *t* = 0.5 s) between naïve and natural geometry for varying *ρ* **c.i**. The estimate of the mean collapses towards zero with increasing *ρ* for naïve geometry, **c.ii**. The estimated variance increases catastrophically with *ρ* for naïve geometry, but only mildly for natural geometry. **c.iii**. Inference accuracy as measured by the mean marginal 2-Wasserstein distance across dimensions decreases for naïve. **d**. Similar analysis when varying *n*_*p*_ for the **d.i**. Mean, **d.ii**. Variance and **d.iii**. mean 2-Wasserstein distance. In panels **c**,**d** shaded error patches show 95% confidence intervals computed via bootstrapping over 100 realizations in all panels; In Figure E.1 we show the full time course of the inferred statistics for one set of network parameters and in Figure E.3 we show the statistics after the full 1.5s, which are qualitatively similar. See Appendix E for detailed numerical methods.

### 3.3 Fast sampling through population geometry using probabilistic spike rules

In the preceding subsection, we have shown how neural population geometry, implemented through neural dynamics, can accelerate the speed of approximate sampling in an efficient balanced network. However, this approach suffers from a fundamental conceptual gap: the firing rates of the spiking network are being used to simulate non-spiking Langevin dynamics; the spiking network itself has not been designed to sample. Specifically, the discretization introduced by the spiking exacerbates the errors introduced by the discrete time implementation of the sampling dynamics, leading to overestimation of the stimulus variance. In this subsection, we take a first step towards bridging this gap by considering an alternative limit of the general model introduced in §2: the case in which sampling is performed leveraging only the stochasticity in the spike rule. Our objective here is not to demonstrate a fully biologically-plausible or practically useful sampling algorithm; rather, it is to illustrate the importance of population geometry in a minimal model for probabilistic spiking.

As in 3.2, we focus on sampling from equicorrelated Gaussian distributions 𝒩 (***μ*, ∑**). In this case, we set **Ψ** = **∑** and ***θ*** = ***μ*** in (10). Naïve sampling here corresponds to choosing **Γ** in some sense generically (see Appendix B.5), while geometry-aware sampling corresponds to choosing **Γ** such that **ΓΓ**^⊤^≃ **∑** up to overall constants of proportionality. In Figures 3, 4, and E.4, we show that naïve choices of **Γ** lead to vanishing spike rates at strong correlations *ρ* and large parameter-space dimensionalities *n*_*p*_. This results in dramatic underestimation of the mean and variance of the target distribution, which is resolved by choosing the geometry appropriately, again allowing inference at behavioral time-scales. Moreover, these networks show Poisson-like variability in spiking statistics (Figure 3), consistent with cortical dynamics [61, 62].

**Figure 3:**
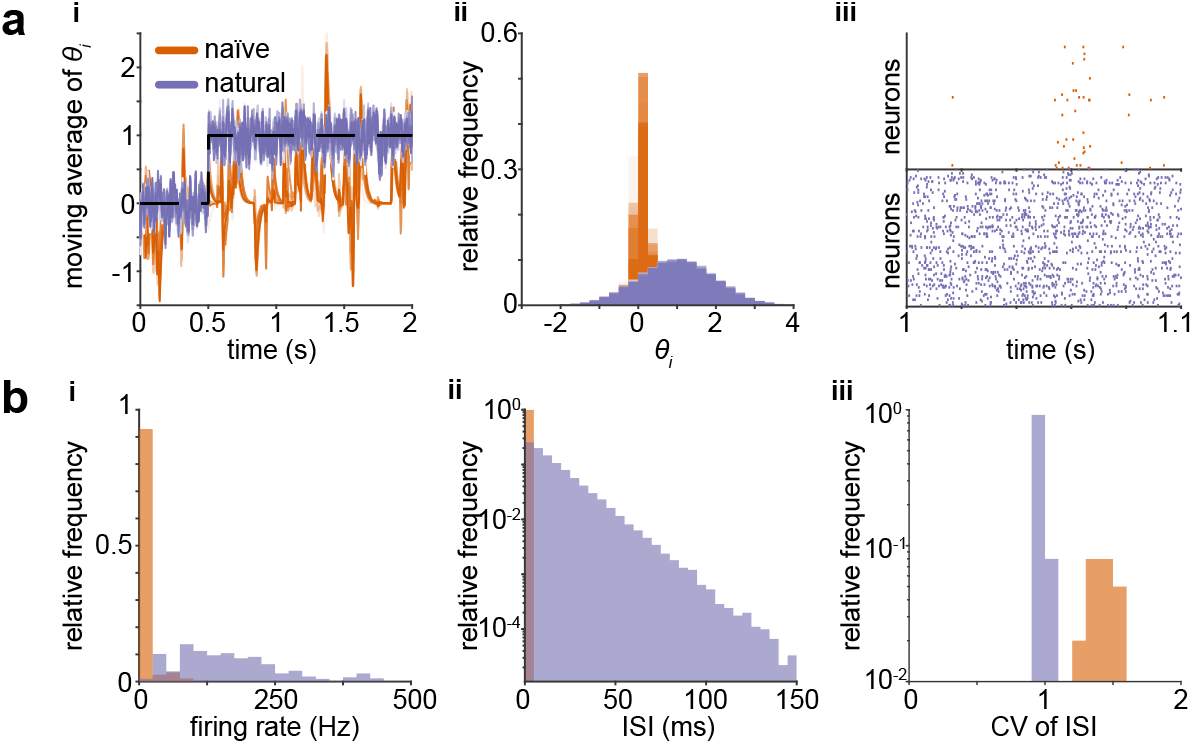
Sampling using probabilistic spiking rules. **a**. Sampling from a *n*_*p*_ = 10-dimensional equicorrelated Gaussian distribution of correlation *ρ* = 0.75 using a network of *n*_*n*_ = 100 neurons. **a.i** 100 ms moving averages of parameter estimates 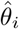 over time, for naïve and natural geometry. At *t* = 0.5 s, the target mean shifts from zero to one, as indicated by the black dashed line. **a.ii**. Marginal distributions of *θ*_*i*_ after stimulus appearance. **a.iii** Spike rasters over a 100 ms window. **b**. Variability in spiking across 1000 trials. **b.i**. Spike rate distributions across neurons and trials for a stimulus distribution as in (a). **b.ii**. The distribution of ISIs for an example neuron across trials is approximately exponential. **b.iii**. The distribution of coefficients of variation of ISIs across neurons. See Appendix E for detailed numerical methods.

**Figure 4:**
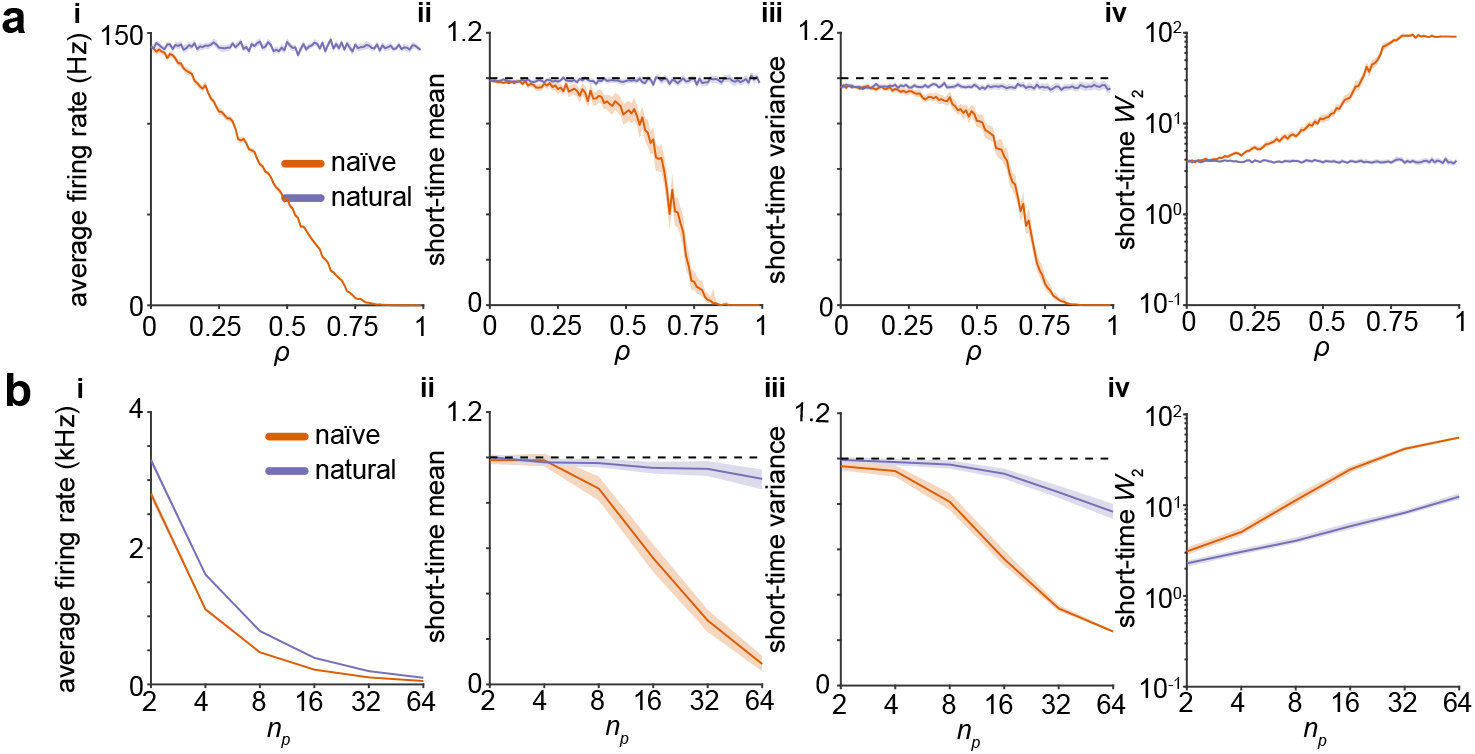
Population geometry enables fast sampling from strongly-correlated high-dimensional distributions in networks with probabilistic spiking rules. **a**. Sampling from strongly-equicorrelated Gaussians in *n*_*p*_ = 10 dimensions using a network of *n*_*n*_ = 100 neurons requires careful choice of geometry. The stimulus setup is as in Figure 3. **a.i**. At strong correlations *ρ*, spiking is suppressed in networks with naïve geometry, but not when the natural geometry is used. **a.ii**. Dimension-averaged estimate of the mean signal in the 50 ms after stimulus onset. See Figure E.4 for steady-state statistics. **a.iii**. As in **ii**, but for the variance. **a.iv**. As in **ii**, but for the mean marginal 2-Wasserstein distance across dimensions. **b**. Sampling in high dimensions requires careful choice of geometry. **b.i**. At moderately strong correlations *ρ* = 0.75 and high dimensions, spiking is suppressed in networks with naïve geometry, but not when the natural geometry is used. Here, we use 10 neurons per parameter, i.e., *n*_*n*_ = 10*n*_*p*_. **b.ii**. Dimension-averaged estimate of the mean signal in the 50 milliseconds after stimulus onset. See Figure E.4 for steady-state statistics. **b.iii**. As in **ii**, but for the variance. **b.iv**. As in **ii**, but for the mean 2-Wasserstein distance. Shaded error patches show 95% confidence intervals computed via bootstrapping over 100 realizations in all panels; see Appendix E for further details.

In Appendix B.5, we provide a more careful analysis of the strongly-correlated limit *ρ* ↑ 1. Informally, we show that the probability of spiking should vanish if **Γ** is chosen sufficiently naïvely and the mean of the target distribution is uniform across parameter dimensions, i.e., 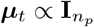. This analysis is not specialized to a particular case, and holds generally for the model of §2. Therefore, careful choice of population geometry, as implemented by the neural code, is required for fast sampling in this model.

## 4 Discussion

In this paper, we have shown how careful choice of neural population geometry enables fast sampling in spiking neural networks. We presented a unified framework in which EBN samplers approximating Langevin dynamics with greedy spiking and approximate Metropolis-Hastings samplers with deter-ministic voltage dynamics and probabilistic spiking can be unified. We then leveraged population geometry to perform rapid sampling at behaviorally-relevant timescales in these two disparate limits of our general model. We now discuss some of the limitations of our work, and highlight possible directions for future inquiry.

Like the original EBN model, the probabilistic spiking model introduced in §2 suffers from the limitations that it requires instantaneous propagation of spike information and that only one neuron is allowed to spike at a time [20, 45–49]. Moreover, the discretization timestep enforces a hard cutoff on the maximum spike rate. Some of these limitations could be partially circumvented by generalizing the spike proposal distribution to allow multiple spikes. However, such a model would still suffer from the issue that an accept/reject step that accounts for the effect of spikes from multiple neurons will require instantaneous communication across the network. This limitation could possibly be overcome within the framework of asynchronous Gibbs sampling [63, 64], which ignores the requirement that updates should be coordinated across the network.

The analysis of §2 shows that the models of §3 can be viewed as limiting cases of a single framework. It is likely that the parallels between these limiting models could be further strengthened by viewing the Gaussian noise in the voltage dynamics of the EBN sampler as an approximation to the effect of the stochastic spiking of other neurons in the Metropolis-Hastings sampler. Such an approximation could be obtained in the limit of large network size given an appropriate treatment of asynchronous updates, provided that one could neglect the coupling between spikes in different neurons induced by the proposal distribution [62, 65–67]. Careful analysis of the relationship between these two sources of stochasticity will be an interesting subject for future investigation. Moreover, it will be interesting to investigate how they might be integrated in a single network, which would result in a spiking sampler somewhat reminiscent of the Metropolis-adjusted Langevin algorithms used in machine learning [68, 69].

Here, we have focused entirely on Gaussian target distributions. As we sketch in Appendices B.8 and C.4, both the probabilistic spiking sampler and the EBN sampler could in principle be extended to general exponential family targets. However, the most naïve extensions to non-Gaussian distributions would involve non-linear and non-local interactions in the voltage dynamics. In particular, one would have to account for state-dependence in the matrix fields **D**(**z**) and **S**(**z**), which would require the inclusion of the field **Φ**(**z**) in the complete recipe (12) (see ref. [43]). Mapping such interactions onto biological mechanisms requires more careful consideration. In recent work, Nardin et al. [70] have proposed how dynamical systems with polynomial nonlinearities can be approximated by deterministic EBNs with multiplicative synapses. This approach could be extended to the stochastic setting of networks designed to sample, which will be interesting to test in future work.

In this work, we did not constrain neurons to follow Dale’s law, and single neurons therefore have both excitatory and inhibitory effects on their neighbors. Many frameworks have been proposed to map the connectivity of unconstrained network algorithms onto distinct excitatory and inhibitory neuron types [9, 71, 72]. These refinements of the biological plausibility will not affect our key argument of accelerating inference through a favorable population geometry. However, different possible implementations that comply with Dale’s law will make different predictions for experimentally-measureable biophysical properties. For example, although less numerous, fast spiking inhibitory neurons have higher firing rates [71, 72], which could allow the approximate symmetry of the readout, as required by the construction of §2, to be maintained.

In biological spiking networks, probabilistic spike emission and probabilistic synaptic release are natural sources of stochasticity [73–76]. These two layers of probabilistic computation provide additional flexibility in processing beyond the simple accept/reject step considered here. As a result, it is likely that one could construct sampling algorithms that are at once more biophysically detailed and more computationally efficient than the simple network constructed in §2. Further investigation of how violations of detailed balance through these mechanisms and the matrix **S** in the “complete recipe” framework could enable faster sampling will be a particularly interesting objective [17, 18, 51]. Moreover, it will be interesting to investigate how natural gradients can accelerate inference in the presence of more complex synaptic dynamics, as studied in recent work by Kreutzer et al. [77].

The algorithms proposed in this work could also enable fast sampling in neuromorphic circuits [78–80]. As they require only the local membrane voltage to compute accept/reject steps, these algorithms could be implemented in a distributed neuromorphic architecture. They would then potentially be limited only by the timescale of individual computing units (which are much faster than biological neurons) rather than the dimensionality of the inference problem [81].

The sampling processes considered in this work focus on short-timescale perceptual inference. However, similar probabilistic inference can occur at longer timescales of learning [82–84]. These long-timescale sampling processes would allow networks to flexibly infer their synaptic weights.

Importantly, learning the task—i.e., learning the matrix **∑**^−1^—and learning the representational geometry—**D, S**, and **Γ**—are distinct processes that can occur in parallel. Once the task structure is learned, adapting the geometry of the population code to match changing stimulus geometry can be achieved through meta-learning [85]. Recent works have analytically studied the population geometry that results from this sampling procedure in rate-based networks [86], and proposed algorithms for efficient learning in EBNs [47, 87–89] and other classes of spiking networks [90–93]. However, the interactions between fast activity sampling and slow network parameter sampling in neural circuits—particularly spiking networks—remain poorly understood [84]. Characterizing how adaptive population geometry accelerates perceptual inference in dynamic environments will be an important step towards a more complete understanding of probabilistic inference in neural circuits.

## Acknowledgments and Disclosure of Funding

PM was supported by the Harvard Mind Brain Behavior Interfaculty Initiative. JAZ-V and CP were supported by a Google Faculty Research Award and NSF Award DMS-2134157. This work was also supported by a grant from the National Institute of Health (R01-DC-017311) to VNM. We thank Tony Zador for co-organizing the NeurIPS 1996 workshop on “Synaptic transmission: reliability and variability”, whose follow-up discussions led to this paper.

## Checklist

1. For all authors…
  a. Do the main claims made in the abstract and introduction accurately reflect the paper’s contributions and scope? [Yes]
  b. Did you describe the limitations of your work? [Yes] We provide a detailed discussion of the limitations of our work in §2 and §4.
  c. Did you discuss any potential negative societal impacts of your work? [N/A] Our work is purely theoretical, and we do not anticipate that it will have negative societal impacts as outlined in the ethics guidelines
  d. Have you read the ethics review guidelines and ensured that your paper conforms to them? [Yes]
2. If you are including theoretical results…
  a. Did you state the full set of assumptions of all theoretical results? [Yes] We state the assumptions of our results in §2 and in Appendices B, C, and D.
  b. Did you include complete proofs of all theoretical results? [Yes] See Appendices B, C, and D.
3. If you ran experiments…
  a. Did you include the code, data, and instructions needed to reproduce the main experimental results (either in the supplemental material or as a URL)? [Yes] Matlab code used to generate all figures is available at https://github.com/Pehlevan-Group/FastSpikingSampler.
  b. Did you specify all the training details (e.g., data splits, hyperparameters, how they were chosen)? [N/A] We did not train any networks
  c. Did you report error bars (e.g., with respect to the random seed after running experiments multiple times)? [Yes] See Appendix E.
  d. Did you include the total amount of compute and the type of resources used (e.g., type of GPUs, internal cluster, or cloud provider)? [Yes] See Appendix E.
4. If you are using existing assets (e.g., code, data, models) or curating/releasing new assets…
  a. If your work uses existing assets, did you cite the creators? [N/A]
  b. Did you mention the license of the assets? [N/A]
  c. Did you include any new assets either in the supplemental material or as a URL? [N/A]
  d. Did you discuss whether and how consent was obtained from people whose data you’re using/curating? [N/A]
  e. Did you discuss whether the data you are using/curating contains personally identifiable information or offensive content? [N/A]
5. If you used crowdsourcing or conducted research with human subjects…
  a. Did you include the full text of instructions given to participants and screenshots, if applicable? [N/A]
  b. Did you describe any potential participant risks, with links to Institutional Review Board (IRB) approvals, if applicable? [N/A]
  c. Did you include the estimated hourly wage paid to participants and the total amount spent on participant compensation? [N/A]

## Supplemental Information

### A Table of variables and parameters used in the models and simulations

We provide a table stating the dimensionality and values taken by the variables and parameters used throughout the paper.

**Table 1.** Variable and parameter names

| Variable name | Description | Value/Size |
| --- | --- | --- |
| $n_p$ | Number of features encoded by the network | 2 – 128 |
| $n_n$ | Number of neurons in the network | 2 – 2560 |
| $n_i$ | Dimensionality of the input signal | usually $n_p$ |
| $\theta$ | Features inferred by the network | $n_p \times 1$ |
| $\Gamma$ | Readout weights | $n_p \times n_n$ |
| $\Psi$ | Covariance of structured Gaussian posterior | $n_p \times n_p$ |
| $\mathbf{V}$ | Membrane voltage | $n_n \times 1$ |
| $\mathbf{T}$ | Spiking thresholds (for each neuron) | $n_n \times 1$ |
| $\Omega$ | Recurrent weights | $n_n \times n_n$ |
| $\tau_m$ | Membrane time constant | scalar; $\sim 20$ ms |
| $\Delta$ | Discretization timestep | scalar |
| $\eta$ | Discrete-time decay constant; $\eta = \Delta/\tau_m$ | scalar |
| $\alpha$ | Elastic net $\ell_1$ penalty | scalar |
| $\lambda$ | Elastic net $\ell_2$ penalty | scalar |
| $\mu$ | Mean of target Gaussian distribution | $n_p \times 1$ |
| $\Sigma$ | Covariance of target Gaussian distribution | $n_p \times n_p$ |
| $\rho$ | Cross-correlation of equicorrelated Gaussian | [0-0.99] |
| $\mathbf{x}$ | Input signal | $n_i$ |
| $\mathbf{A}$ | Input weights | $n_p \times n_i$ |
| $\mathbf{W}$ | Standard Brownian motion | usually $n_p \times 1$ |

### B Approximate Metropolis-Hastings sampling using probabilistic spiking rules

In this Appendix, we provide a step-by-step construction of the spiking sampler introduced in §2 of the main text. As introduced in §2 of the main text, our goal is to use

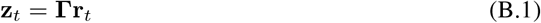

to sample a Gaussian distribution *P* (**z**) with mean ***θ*** and covariance **Ψ**, where

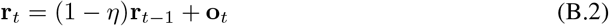

is the filtered spike history for a decay constant 0≤ *η* ≤ 1 (note that 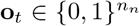). In §B.1, we construct a circuit that samples a Gaussian distribution using a discrete-time perfect integrator of the spike train (i.e., *η* = 0). In §B.2, we relax this assumption, yielding an approximate sampler using leaky integration in discrete-time, and discuss the behavior of this model in the continuum limit. In general, the proposal distribution could be computed using a stochastic gradient step followed by an accept/reject step, yielding Metropolis-adjusted Langevin dynamics [68, 69]. In this case, the accept/reject step allows the algorithm to compensate for some of the sampling error introduced by discretization at the expense of needing to compute a likelihood ratio, which can be expensive for high-dimensional distributions.

#### B.1 A simple sampler assuming perfect integration and balance

We first construct a sampling circuit under the assumptions that we have access to a perfect integrator of the spike train, i.e. that *η* = 0, and that the readout matrix is of the form

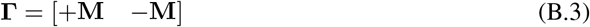

for some matrix 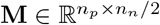.

At the *t*-th timestep, we choose one neuron *j* uniformly at random, and let the spike proposal be **o**^*′*^ = **e**_*j*_, where **e**_*j*_ is the *j*-th standard Euclidean basis vector (i.e., (*e*_*j*_)_*k*_ = *δ*_*jk*_). This yields a candidate readout

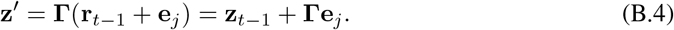

Under the symmetry assumption on **Γ**, the acceptance ratio is given by

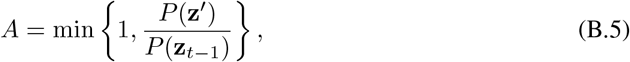

as the proposal distribution is exactly symmetric in **z**^*′*^ and **z**_*t*_. Then, we accept the proposed spike with probability *A*. Concretely, for *u* ∼ *𝒰* [0, 1], we take

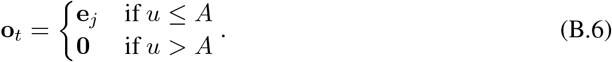

For *P* (**z**) a Gaussian distribution with mean ***θ*** and covariance **Ψ**, we have

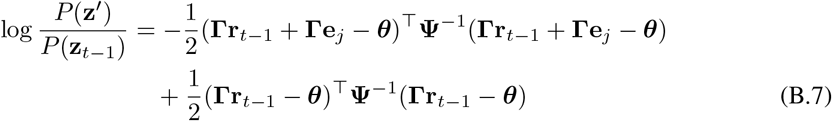

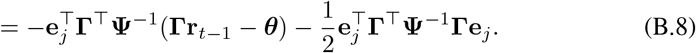

Defining the matrix

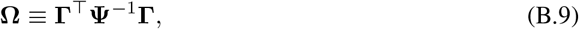

we interpret the first term in the log-odds ratio as a membrane potential,

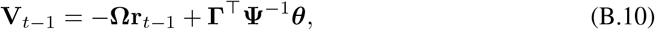

and the second as a threshold

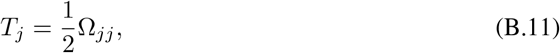

such that the acceptance ratio is

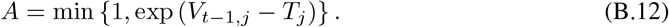

With this definition, the membrane voltage simply integrates the spike train:

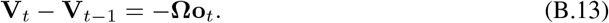

We now observe that, if *n*_*n*_ > *n*_*p*_, it is possible that a spike may not contribute to the parameter estimate. Concretely, we say that a spike, or the corresponding neuron, is *irrelevant* if it is annihilated by **Γ**, i.e., if **Γe**_*j*_ = **0**. Irrelevant spike proposals are always accepted with probability one, as we have **z**^*′*^ = **z**_*t*− 1_. Moreover, the membrane voltage is not changed by the emission of irrelevant spikes. For these reasons, we are free to re-define the population rate **r**_*t*_ to exclude such spikes. Therefore, a timestep with an irrelevant spike is equivalent to not updating the network at all, and we could choose to re-define the network such that only relevant neurons are included. However, though there will exist some non-trivial set of vectors that are annihilated by **Γ** if *n*_*n*_ > *n*_*p*_, the situation in which this null space is axis-aligned (thus implying the existence of irrelevant neurons) is not generic.

We note that appropriate initialization of the membrane potential (for the desired mean) is important, as otherwise some bias will be introduced. Thus, as written, this model cannot easily accommodate a time-varying mean signal ***θ***_*t*_. This shortcoming could obviously be addressed by taking

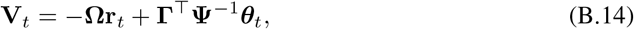

which yields voltage dynamics

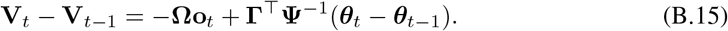

Alternatively, one could also take

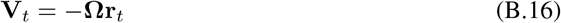

and re-define the threshold for the *j*-th neuron to be time-varying:

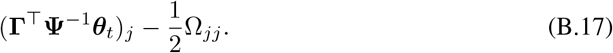

The former of these approaches is reasonable from a biological perspective, as the new term **Γ**^⊤^**Ψ**^−1^(***θ***_*t*_− ***θ***_*t*−1_) in the voltage dynamics has the interpretation of a signal ***θ***_*t*_ −***θ***_*t*−1_ fed through an input weight matrix **Γ**^⊤^**Ψ**^−1^. For this sampling procedure to work, the mean should be slowly-varying.

#### B.2 Relaxing the assumption of perfect integration

We now relax the assumption of the perfect integration, and assume only that *η* ≪ 1. With the same proposal distribution as before, we take the acceptance ratio of the accept-reject step to be

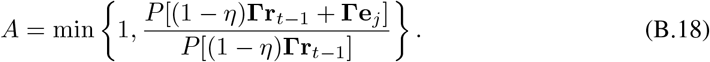

As noted in the main text, this choice implements a sort of look-ahead step. Moreover, if we took the acceptance ratio to depend on the likelihood of the proposed state with decay, *P* [(1 −*η*)**Γr**_*t*− 1_ + **Γe**_*j*_], relative to the likelihood *P* [**Γr**_*t*− 1_] of the current state (rather than the next state with decay but without the proposed spike), the resulting log-odds ratio would include terms of order *η* that are quadratic in the rate **r**_*t*_.

With the choices above, the log-odds ratio is

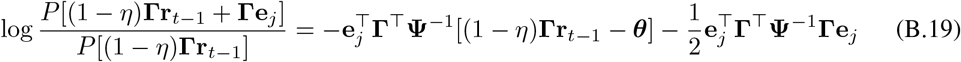

which, like in the perfect integrator model, is linear in the rate. As in the perfect integrator model, we define the recurrent weight matrix

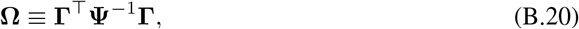

and interpret the first term in the log-odds ratio as a membrane potential,

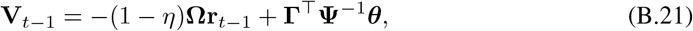

and the second as a threshold

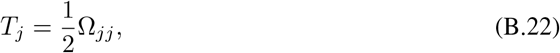

such that the acceptance ratio is

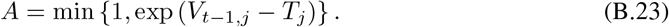

With this definition, the membrane voltage evolves as

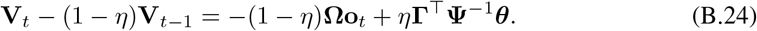

Therefore, this model differs from the perfect integrator model of §B.1 only in the voltage dynamics; the perfect integrator is recovered exactly if we set *η* = 0. As in the perfect integrator model, the natural generalization of these leaky dynamics to time-varying mean signal ***θ***_*t*_ is to take

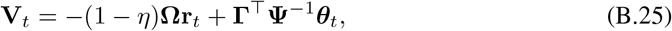

which leads to the dynamics

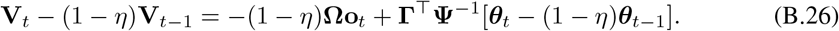

This will, of course, not be an exact sampler unless the mean is constant. If the mean is slowly-varying, however, it should be a good approximate sampler. We remark that we can re-write the membrane voltage as

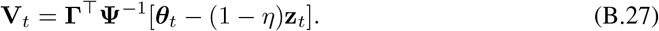

#### B.3 Adding an elastic net prior on the rates

We now consider the effect of adding an elastic net prior on the rates, as was considered in the original work of Boerlin et al. [45]:

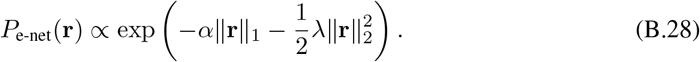

Without loss of generality, we consider the case in which a decay term is included, as we can then recover the perfect integrator by setting *η* = 0. Again defining the acceptance ratio in terms of a comparison against the rate with decay but without the proposed spike, the addition of the prior adds

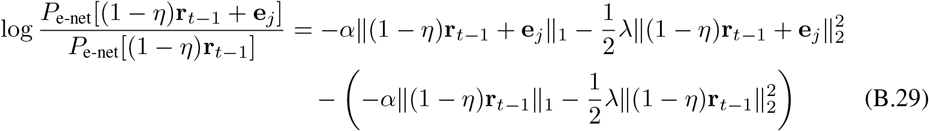

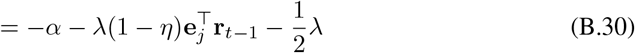

to the log-odds ratio. This yields a modified recurrent weight matrix

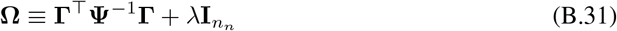

and a modified threshold

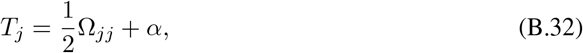

but the expression for the membrane voltage in terms of these parameters is identical in functional form:

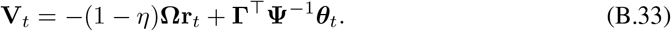

Therefore, adding the elastic net prior changes the definitions of the weight matrix that maps rates to voltages and of the threshold, but not the overall form of the result, hence it does not introduce any new conceptual difficulties. We remark that we could include the constant factor *α* in the membrane voltage as we do in our implementation of EBNs (see Appendix C) rather than in the threshold, which would give it a somewhat different biological interpretation but would have no algorithmic effect.

#### B.4 The continuous-time limit

In this subsection, we consider the continuous-time limit of the models introduced above. This limit corresponds to taking the limit in which spike proposals are made infinitely often, and regarding the dynamics written down previously as a forward Euler discretization of an underlying continuous-time system. For clarity, we write the discrete timesteps, denoted in previous sections simply by *t*, as *t*_*d*_ here, and reserve the unsubscripted symbol *t* for the continuum variable. For a timestep Δ, we let *t* = Δ*t*_*d*_, and let the discrete-time decay rate be *η* = Δ*/𝒯*_*m*_ for a ‘membrane’ time constant *𝒯*_*m*_. We may then write the discretized rate dynamics (B.2) as

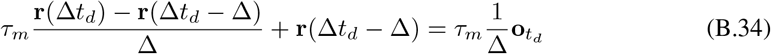

which, taking Δ ↓ 0, of course yields the familiar continuous-time dynamics

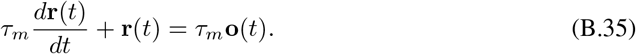

In continuous time, the spike train **o**(*t*) is now composed of Dirac delta functions, as the discretized spikes are rectangular pulses of width Δ in time and height 1*/*Δ. We next consider the voltage dynamics of the leaky integrator for a varying mean signal (B.26), which may similarly be re-written as

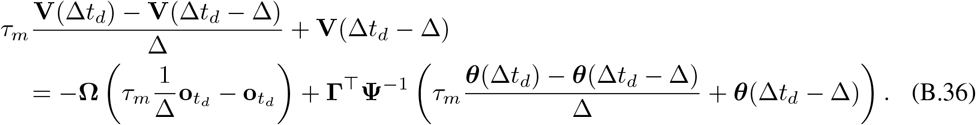

In the continuum limit, we retain only the contribution of the first of the two terms involving the discrete-time spike train, as the other yields pulses of width Δ and height unity, which yield a negligible contribution to the integral. Thus, we have the dynamics

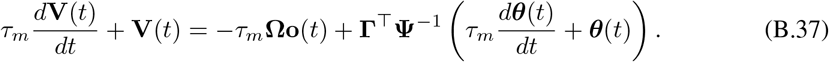

From these dynamics, one could then recover the continuum limit of the perfect integrator dynamics by taking *𝒯*_*m*_ → ∞. In this limit, the rate will decay by only a infinitesimal amount between a rejected spike proposal and the next proposal, meaning that the error incurred by neglecting the asymmetry in the proposal distribution due to the decay should be negligible.

### B.5 Analyzing the strongly-correlated limit of sampling from an equicorrelated Gaussian

Here, we consider how these models behave when sampling from an equicorrelated Gaussian distribution, i.e., a distribution with covariance matrix

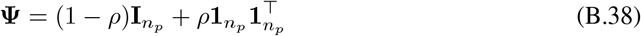

for correlation coefficient *ρ*∈ (−1, +1). We are particularly interested in the strongly-correlated limit *ρ* ↑ 1. As we will consider the case in which the marginal variance does not scale with the correlation, our choice of unit marginal variance is made without loss of generality. For this covariance matrix, the Sherman-Morrison formula yields [94]

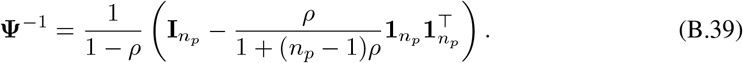

We first consider the case in which the mean signal ***θ*** is identically equal to zero. We argue that, for choices of **Γ** that are in some sense sufficiently naïve, the probability that relevant spikes are emitted should tend to zero as *ρ* ↑1. This corresponds to showing that *V*_*t,j*_ −*T*_*j*_→ −∞ as *ρ* ↑1 for all indices *j* corresponding to relevant neurons. For ***θ*** = **0**, we have

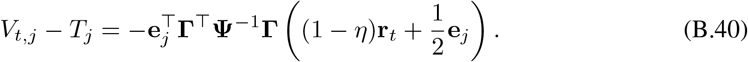

Consider the first timestep, with **r**_0_ = **0**, for which we have

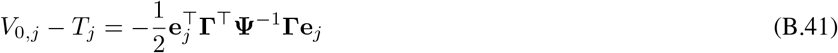

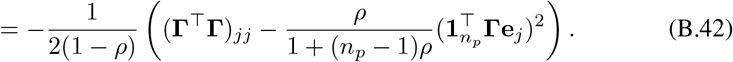

Near *ρ* = 1, we then have the expansion

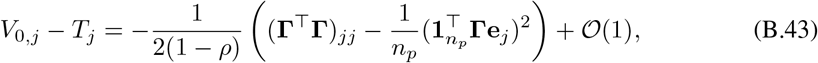

under the assumption that **Γ** is an (1) function of *ρ* in this region, and thus cannot introduce additional possible divergences. For relevant spikes, we have the strict inequality (**Γ**^⊤^**Γ**)_*jj*_ *>* 0. Moreover, by the Cauchy-Schwarz inequality, we have 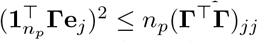, with equality if and only if 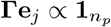. If 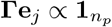, then spikes in neuron *j* affect the readout precisely only along the common mode, and *V*_0,*j*_ −*T*_*j*_ does not diverge as *ρ* ↑ 1. However, this case is quite fine-tuned. For generic **Γ** not satisfying this alignment condition, we have the strict inequality

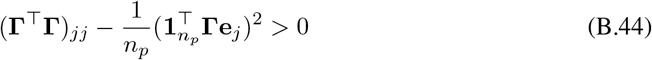

for all relevant neurons. Then, if (**Γ**^⊤^**Γ**)_*jj*_ vanishes no more rapidly than 1 ↑*ρ* as *ρ* 1—i.e., if (**Γ**^⊤^**Γ**)_*jj*_*/*(1− *ρ*) → ∞ as *ρ* 1—we have that *V*_0,*j*_ −*T*_*j*_→ −∞ as *ρ* ↑ 1. This holds, for instance, if (**Γ**^⊤^**Γ**)_*jj*_ is a constant function of *ρ*. Thus, under these conditions, the probability that a relevant spike is emitted at the first timestep vanishes as *ρ* ↑ 1. Heuristically, this in turn implies that the (relevant component of the) rate at the second timestep will be zero with probability one. Therefore, we may iterate this argument forward in time, showing that the probability of emission of relevant spikes should vanish in the limit *ρ* ↑ 1. This argument will not be affected by the addition of an elastic net penalty unless the coefficients *α* and *λ* are taken to diverge as *ρ* is taken to unity, as the coefficients are strictly non-positive.

The situation is somewhat more complicated if the mean signal is not identically zero, in which case we have

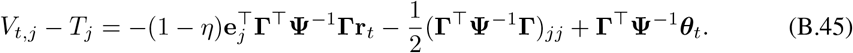

Following our previous analysis, at the first timestep we have

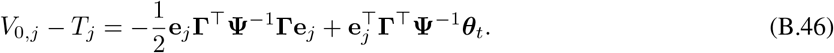

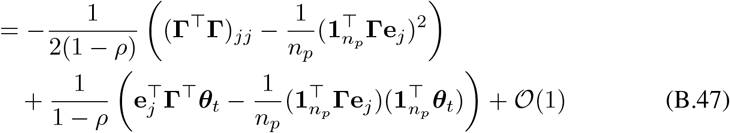

near *ρ* = 1. There are now two possible divergent terms, which can compete to change the sign of *V*_0,*j*_ − *T*_*j*_ as *ρ* ↑ 1. One case of interest is when 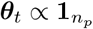. Then,

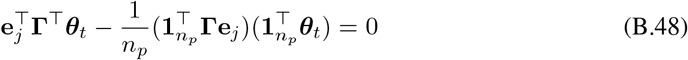

and we have

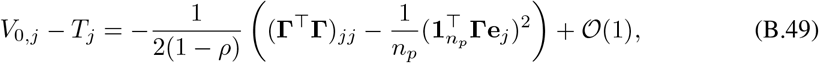

as in the case when ***θ*** was strictly zero, implying that *V*_0,*j*_ −*T*_*j*_→ −∞ under the abovementioned conditions on **Γ**. Another illustrative case is ***θ***_*t*_ = **Γe**_*j*_. With this fine-tuning,

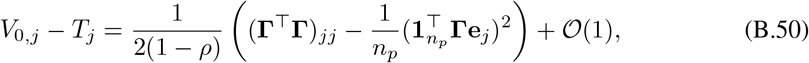

hence we expect *V*_0,*j*_− *T*_*j*_ →+∞ as *ρ* ↑1 under the abovementioned constraints on **Γ**. Thus, in this case, the spike probability should tend to one, showing that complications can arise in the case of a non-uniform mean signal.

### B.6 Analyzing the small- and large-variance limits

We now consider how these models behave in the limits in which the variance of the target distribution is small or large, or, nearly equivalently, the limits in which the scale of the readout matrix is large or small, respectively. In §2.3 of the main text, we argued that the EBN is recovered in the limit in which the variance of the target distribution tends to zero. Concretely, we let 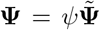 for some fixed matrix 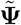, and take the zero-variance limit *ψ* ↓ 0. In terms of the renormalized voltage 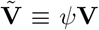, weight matrix 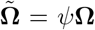, and threshold 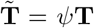, the spike acceptance ratio is 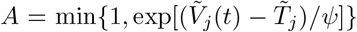. This tends to 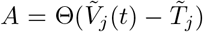 as *ψ*↓ 0 assuming that the re-scaled variables remain order one, recovering the greedy spiking rule used in the EBN. If we instead take the large-variance limit *ψ* ↑ ∞, the acceptance ratio tends to 1 under the assumption that 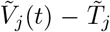 remains 𝒪 (1). Then, each spike proposal is accepted with probability one, and the total firing rate of the population is 1*/*Δ spikes per second for a timestep Δ, meaning that the population-averaged rate is 1*/*(*n*_*n*_Δ) spikes per second.

Re-scaling the readout matrix **Γ** differs from re-scaling the target covariance matrix **Ψ** because the target mean and recurrent spikes appear with the same power of **Ψ** (in particular, **Ψ**^−1^) but different powers of **Γ** in the voltage dynamics, scaling as 𝒪 (**Γ**) and 𝒪 (**Γ**^2^), respectively. Thus, in the large-**Γ** limit in discrete time, the contribution of the target mean should be negligible relative to that of the recurrent spiking input, while the opposite should hold in the small-**Γ** limit. However, these different scalings should not affect the limiting behavior of the acceptance ratio. In Figure B.1, we probe how sweeping the scale of **Γ** over five orders of magnitude affects sampling from a Gaussian distribution of fixed variance. We show that the population-averaged spike rate tends to 1*/*(*n*_*n*_Δ) spikes per second in the small-**Γ** limit, while the spike rate tends to zero in the large-**Γ** limit.

**Figure B.1.**
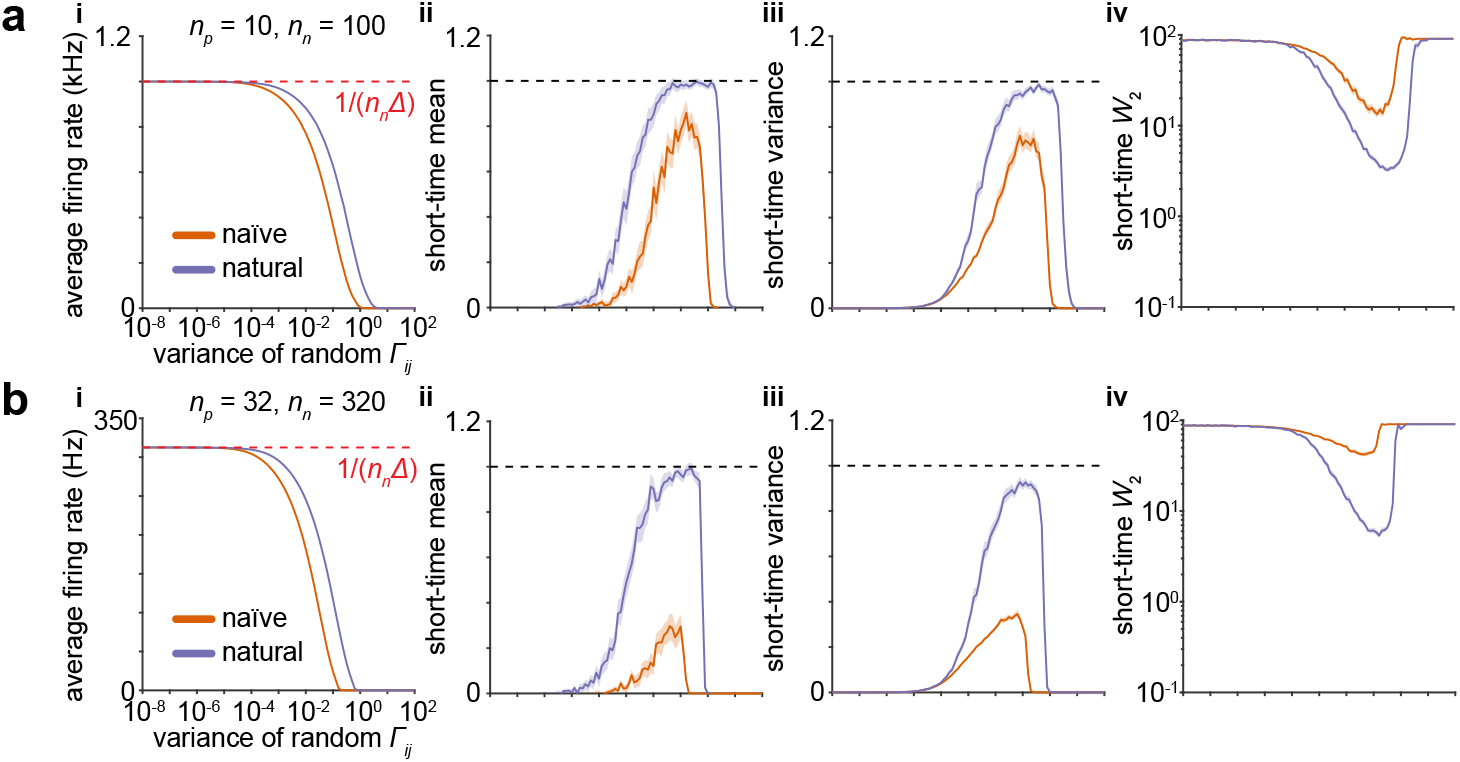
Sensitivity of the Metropolis-Hastings sampler to the scale of the readout matrix. **a**. Sampling performance of a network of *n*_*n*_ = 100 neurons sampling from a *n*_*p*_ = 10-dimensional equicorrelated Gaussian with *ρ* = 0.75 depends on the variance of the elements of the random readout matrix **Γ**. The stimulus setup is as in Figure 3. **a.i**. At large variance, spiking is suppressed in networks with naïve or natural geometry. At small variance, spikes are accepted with probability one, and the population-averaged firing rate tends to 1*/*(*n*_*n*_Δ), as indicated by the red dashed line. **a.ii** Dimension-averaged estimate of the mean signal over the entire stimulus interval. **a.iii**. As in **ii**, but for the variance. **a.iv**. As in **ii**, but for the 2-Wasserstein distance. **b**. As in **a**, but for a network of *n*_*n*_ = 320 neurons sampling from an *n*_*p*_ = 32-dimensional Gaussian. Shaded error patches show 95% confidence intervals computed via bootstrapping over 100 realizations in all panels; see Appendix E for further details.

### B.7 Alternative spike proposal distributions and membrane voltage bounds

In this appendix, we have considered the simplest possible spike proposal distribution: at each timestep, we choose one neuron uniformly at random as a candidate. With this choice, obviously, only one neuron can spike at each timestep, which parallels the spiking rule used in the EBN: only the neuron with the maximum membrane voltage among the entire population is allowed to spike [20, 45]. With a discretization timestep Δ, the maximum total spike rate is then 1*/*Δ. In the EBN literature, previous analysis has shown that this constraint is vital to avoid pathological spiking patterns [20, 45–48], and introduced alternatives such as Poisson spiking rules [48] or hand-tuned refractory periods [49]. We remark that, in these models, the membrane voltage is always strictly bounded from above, which does not hold in our setting. As reset is achieved only through spike emission, and is strictly speaking a decrement of the voltage rather than a true reset, a neuron can exceed the threshold voltage if it is not chosen as a candidate to spike.

### B.8 Sampling from non-Gaussian exponential families

Though our main focus is on Gaussian target distributions, in this appendix we briefly discuss the possibility of constructing a probabilistic spiking sampler for other exponential families. For a distribution with density

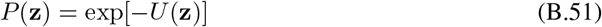

for an energy function *U*, the acceptance ratio (B.18) becomes

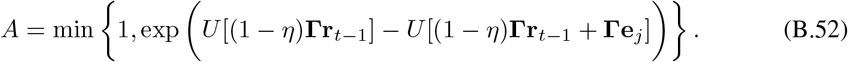

In analogy to the Gaussian case, one could then define the difference of energies to be the difference of the membrane voltage and the threshold. However, the resulting membrane voltage would be a nonlinear function of the firing rate, and would in general evolve according to non-linear dynamics.Most directly, assuming ∥**Γe**_*j*_∥ to be small and *U* to be not too quickly varying, one could make the second-order approximation

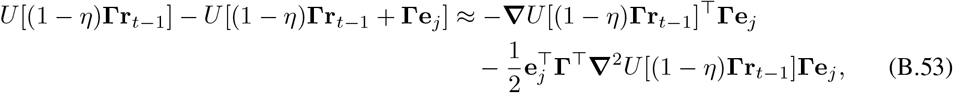

and define the membrane voltage and threshold as

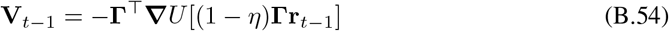

and

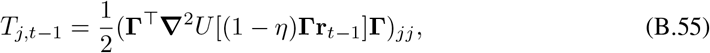

respectively. With this choice, the threshold would be state-dependent, but one could incorporate its state-dependence into a re-defined voltage.

## C. Sampling in efficient balanced networks

### C.1 Encoding a dynamical system in efficient balanced networks

In this section, we provide a pedagogical derivation of the efficient balanced network [20, 45] using the notation we use throughout the paper. The goal of the network is to encode an estimate **z**(*t*) of a signal ***θ*** (a vector of size *n*_*p*_*×* 1 in a population of *n*_*n*_ spiking neurons. The estimate is obtained by reading out a low pass version of the population spiking activity:

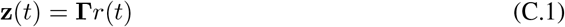

where **Γ** is the *n*_*p*_ *× n*_*n*_ readout matrix and *r*(*t*) is the low-pass filter spike history:

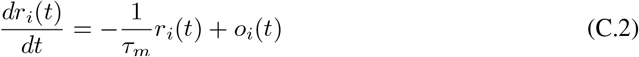

where *𝒯*_*m*_ is the time constant of the readout neurons and *o*_*i*_ is the spike train of neuron *i, o*_*i*_ = 1 if the neuron spiked at *t* and *o*_*i*_ = 0 otherwise.

The goal of the network is to minimize the squared error between the signal and the estimate with an elastic net prior on the firing rate in order to find a good solution while keeping the population spiking activity relatively low:

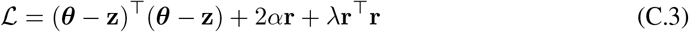

In the standard implementation of efficient balanced networks, neurons use a greedy spiking rule. A neuron should fire if emitting a spike will lower the loss function, i.e., if

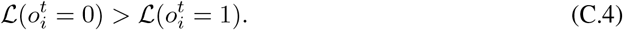

Using the loss function and the definition of the estimate (**z**(*t*) = **Γr**(*t*)), we can rewrite the spiking rule for neuron *i* as:

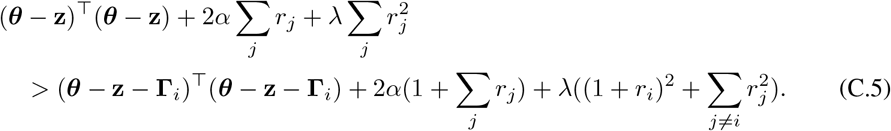

Removing terms that appear on both sides, we get

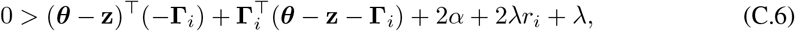

which we can further simplify to

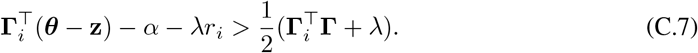

The term on the left hand can be interpreted as the voltage potential of neuron *i*:

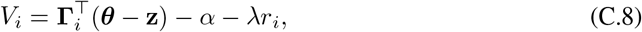

while the term on the right hand size can be interpreted as the firing threshold^5^:

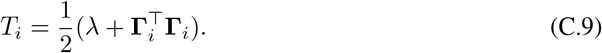

If the voltage potential exceeds this threshold, then the neuron will fire and lower its voltage potential back below threshold. Using (C.8) we can express the dynamics of the voltage potential as a function of the dynamics of ***θ*** and its estimate **z**:

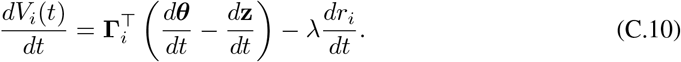

We can rewrite this equation to obtain an expression for the membrane dynamics as a function of the signal, the firing rates, and the spike trains of the neurons. By adding and subtracting 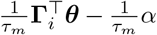 and noting that 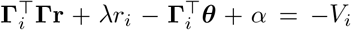, we further simplify the expression to obtain membrane dynamics as a function of the membrane potential, the effect of a new spike on the circuit and the encoded dynamical system:

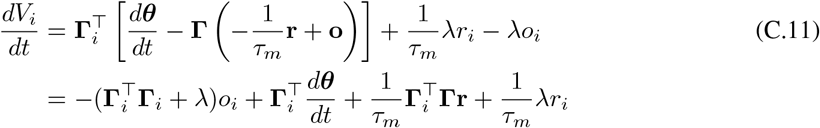

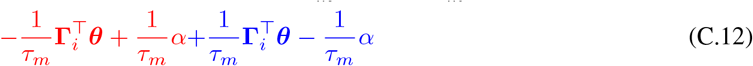

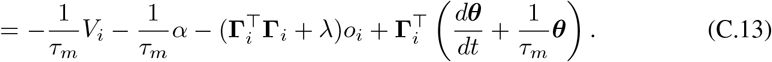

We therefore obtain the membrane dynamics for the efficient balanced spiking network proposed in [20, 45]. We can rewrite the dynamics in vector form:

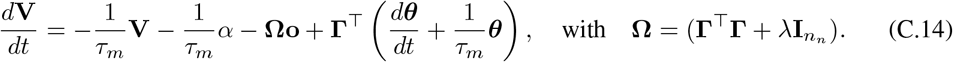

Using this scheme one can encode of variety of dynamical systems, including Langevin dynamics [20, 46, 47].

### C.2 Sampling in efficient balanced networks using naïve Langevin dynamics

Using the scheme presented in §C.1 above, Savin and Denève [20] proposed to implement a dynamical system corresponding to the naïve Langevin dynamics of a multivariate normal distribution. We use a linear Gaussian model similarly to several studies of neuroscience inspired sampling-based networks [9, 20]. These networks estimate the posterior probability of hidden sources (***θ***) given sensory inputs **x** corrupted by Gaussian noise, 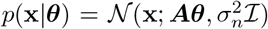, and prior expectations on the values of the hidden sources *p*(***θ***) = 𝒩 (***θ***; 0, ***C***). The mean ***μ*** and covariance **∑** of the posterior probability of the features given the input, *p*(***θ***|**x**) ∝ *p*(**x**|***θ***)*p*(***θ***), are 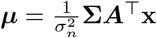 and 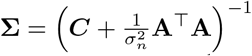 respectively. Up to an irrelevant constant offset, the corresponding energy function is 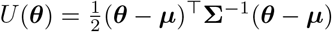, and its gradient is Δ*U* (***θ***) = **∑**^−1^(***θ*** − ***μ***). We define *𝒯*_*s*_ as the timescale of the inference process and we set 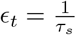. Then, we can write down the naïve Langevin dynamics from (11):

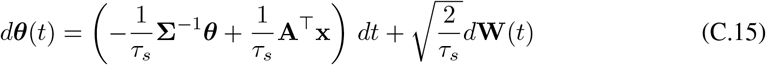

These dynamics can be approximated by an efficient balanced network by replacing in (C.14) ***θ*** by **z**. As discussed in [20], this approximation will introduce an acceptable error in most practical situations:

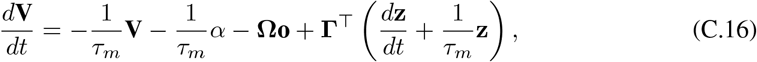

and we therefore obtain the membrane dynamics for the efficient balanced network proposed by Savin and Denève [20]:

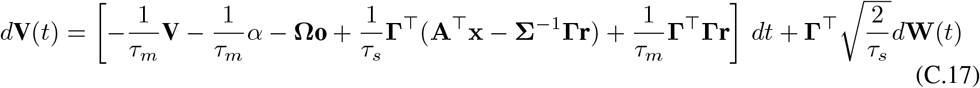

Note here that we have two timescales: the timescale of neuronal representations *𝒯*_*m*_ ∼20 ms, controlled by the biophysical properties of the neurons, and the timescale *𝒯*_*s*_ of the Langevin diffusion encoded by the network.

### C.3 Sampling in efficient balanced networks using the complete recipe for stochastic gradient MCMC

The model proposed by Savin and Denève [20] implements naïve Langevin dynamics, which are known to be slow in high dimensions. Instead, we can use the “complete recipe” for stochastic gradient MCMC [43] to write another sampler with the same equilibrium distribution but more favorable convergence properties. For any positive semi-definite matrix **D** and skew-symmetric matrix **S**, the following dynamics will converge to *N* (***μ*, ∑**) as their stationary distribution:

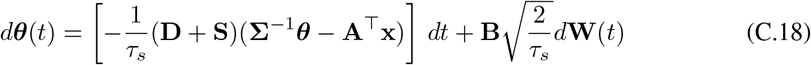

with **BB**^⊤^ = **D**.

We can use (C.14) to encode this more general formulation into an efficient balanced network, yielding the following membrane dynamics:

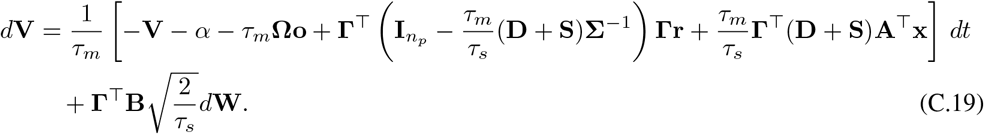

Any choice of positive semi-definite matrix **D** leads to a valid sampler, but extensive work inspired by Amari’s seminal work on natural gradient descent [39, 40] has shown that a principled choice is to take **D** to be the inverse of the Fisher information matrix **G** [42, 43]. For the multivariate Gaussian distribution, the Fisher information matrix is given by:

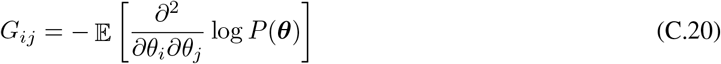

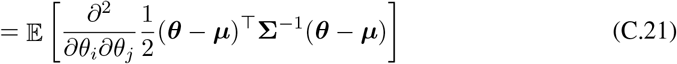

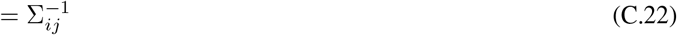

We should therefore choose **D** = **G**^−1^ = **∑**. Note that for the multivariate Gaussian distribution, the Fisher information matrix is identical to the Hessian and is location independent - it does not depend on the value of ***θ***. For more complex distributions, the Fisher information matrix might be difficult to compute and an approximation can be used as long as it is valid (positive semi-definite) within the complete recipe framework [43, 58, 95] and state dependent matrices can be corrected for using the term **Φ** from the complete recipe in (12) [43].

In this work, we have considered only hand-tuned or random choices for the matrices controlling the geometry. Previous work by Hennequin et al. [9], when framed within the complete recipe, proposes methods to find a skew-symmetric matrix **S** which accelerates the dynamics. Non-reversibility is indeed known to accelerate learning, but analysing networks with such dynamics is notoriously difficult [28, 96]. In contrast, approximations for the inverse Fisher information matrix are readily computable, even in non-Gaussian settings [58, 95].

### C.4 Sampling from non-Gaussian exponential families using efficient balanced networks

To sample from a non-Gaussian exponential family distribution with energy *U* (***θ***) using an EBN, we can simulate the “complete recipe” for a general exponential-family as stated in (12) of the main text, which we reproduce below:

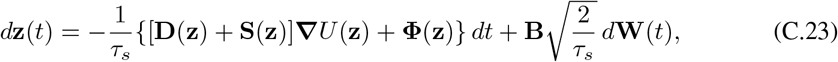

where **BB**^⊤^ = **D** [13]. Here, we have introduced an auxiliary time constant *𝒯*_*s*_, as in our previous discussion of the Gaussian case. Then, the EBN voltage dynamics (C.14) become

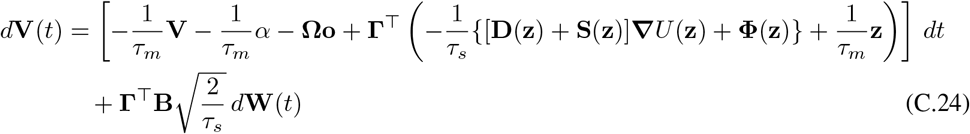

where we have again made the approximation of the Langevin sampling trajectory by the approximate sampling trajectory **z** =. It is easy to see that these dynamics will in general involve non-linear dependence on the population firing rate **r** through the readout **z** = **Γr**.

## D Natural gradient enables fast sampling in linear rate networks

In this section, we analyze how natural gradients enable fast sampling in rate networks designed to sample from zero-mean Gaussian distributions. Our starting point is the complete recipe for sampling from a zero-mean Gaussian distribution with covariance **∑**:

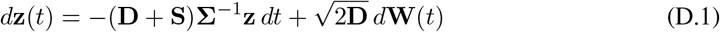

where **D** is a symmetric positive-semidefinite matrix and **S** is skew-symmetric. In previous work, Hennequin et al. [9] showed how these dynamics can be interpreted as a linear rate network, and demonstrated that careful choice of **S** can accelerate sample autocorrelation timescales in networks with 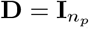. Here, our objective is to show how the geometry of inference, set by **D**, can accelerate sampling in these rate networks.

Our analysis is a straightforward application of the classic theory of Ornstein-Uhlenbeck processes [29, 52, 97]. Assuming for simplicity a deterministic initial condition **z**(0) = **0**, the solution to this stochastic differential equation is given by the Itô integral

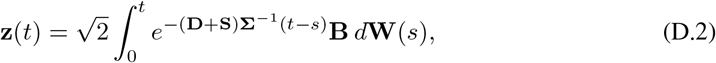

which has ensemble mean zero and ensemble covariance

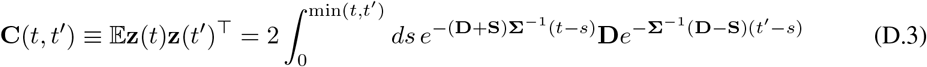

The ensemble distribution of **z**(*t*) is of course Gaussian, with mean zero and covariance given by the equal-time covariance function **C**(*t*) ≡ **C**(*t, t*).

We assume that (**D** + **S**)**∑**^−1^ is a non-defective matrix with all eigenvalues having positive real part, such that the process has a well-defined stationary state. By construction, the covariance matrix of the stationary state is precisely the target covariance matrix **∑**. In the stationary state, we have a simplified two-point function

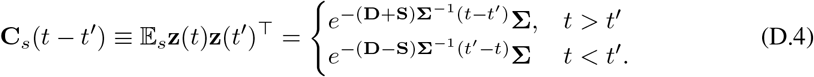

where we have observed that

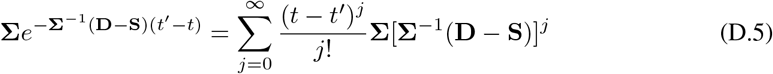

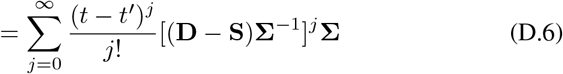

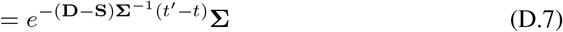

One standard case in which the integral defining the non-stationary covariance function can be evaluated explicitly is if **A** = (**D** + **S**)**∑**^−1^ is a normal matrix (i.e., if **AA**^⊤^ = **A**^⊤^**A**) [29]. Then, there exists a unitary matrix **U** such that 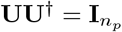 and

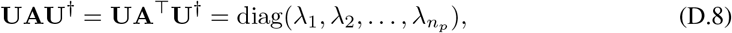

which, for all *t* ≥ *t*^*′*^, yields

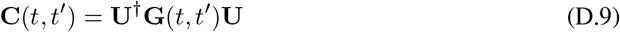

for

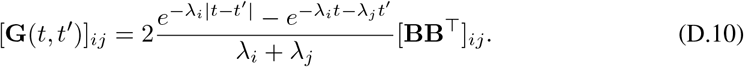

We will measure the rate of convergence of the ensemble distribution to the stationary distribution in the Kullback-Leibler (KL) divergence and 2-Wasserstein distance for several choices of **D**. We note that this is not the same measure as considered in our numerical simulations, where we examine the distribution of samples over time within a single trajectory, and measure the mean 2-Wasserstein distance between univariate marginals. As noted in Appendix E, an organism must usually make estimates based on the distribution of samples over a single trajectory. However, even at equilibrium, it is challenging to analytically characterize the 2-Wasserstein distance between samples from a Gaussian distribution and the underlying population distribution [60]. Therefore, we will consider the ensemble distribution at a given time. We denote the KL divergence and 2-Wasserstein distances as a function of time by *K*(*t*) and *W*_2_(*t*), respectively. As all distributions of interest are Gaussian, we have the relatively simple formulas

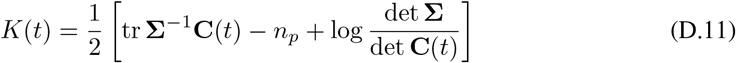

and

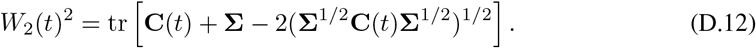

### D.1 Naïve Langevin dynamics

We first consider naïve Langevin sampling with 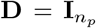 and **S** = **0**. In this case, **A** = (**D** + **S**)**∑**^−1^ = **∑**^−1^ is symmetric and therefore normal, hence, letting the diagonalization of **∑** be

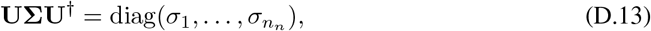

the result above yields

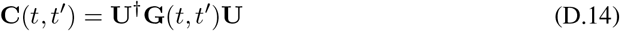

for

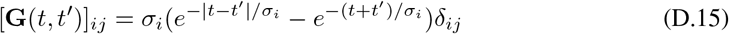

for all *t* ≥ *t*^*′*^. In particular, equal-time covariances are governed by

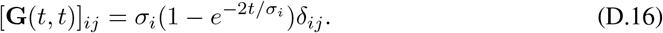

In matrix form,

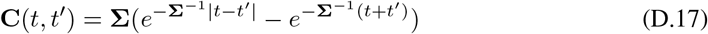

and

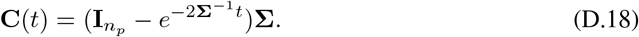

This choice yields stationary covariance

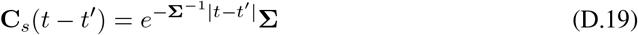

hence large eigenvalues of **∑** will introduce long autocorrelation timescales.

In this case, *K*(*t*) and *W*_2_(*t*) are simple to compute thanks to the fact that **∑** commutes with **C**(*t*), yielding

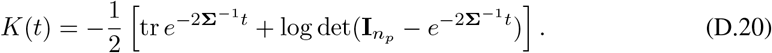

and

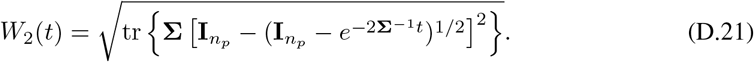

In terms of the eigenvalues *s*_1_, …, *s*_*n*_ of **∑**, these distances are

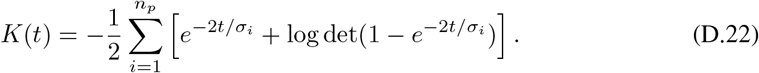

and

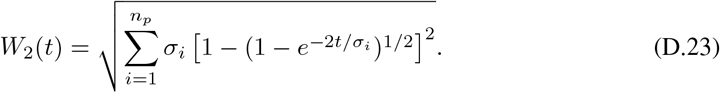

### D.2 Sampling in the space of natural parameters

We now consider sampling in the space of natural parameters, with **D** = **∑** and **S** = **0**. In this case, 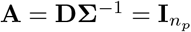 is trivially normal, hence, for *t* ≥ *t*^*′*^,

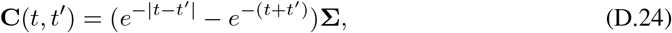

with

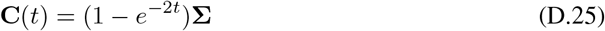

in particular. In this case, the stationary covariance is

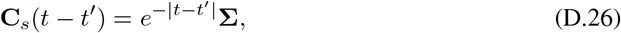

hence the stationary autocorrelation timescale will be independent of the spectrum of **∑**.

This setup can also easily be generalized to the case in which **S***?*= **0**. In this case, 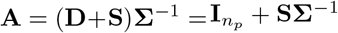 is not in general a normal matrix. However, we can evaluate the covariance function by exploiting the particular structure of the problem. Factoring out the terms involving the identity matrix and using the skew-symmetry of **S**, we have

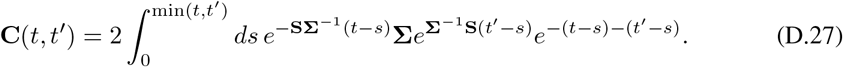

We now observe that

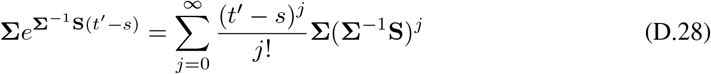

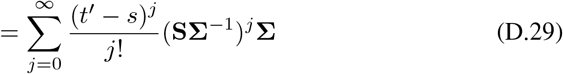

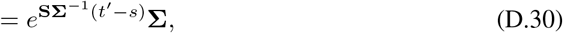

hence 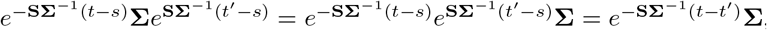, and therefore

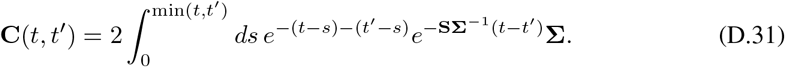

The equal-time covariance is then

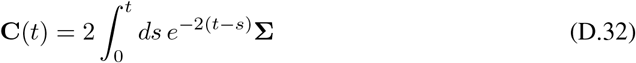

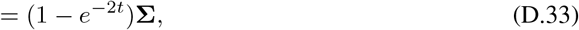

as in the case **S** = **0**. Thus, the rate of convergence of the ensemble sampling distribution to the stationary distribution will be unaffected if we add this skew-symmetric term. With the addition of the skew-symmetric term, the stationary covariance is

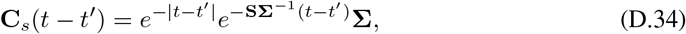

which reflects the fact that this term introduces non-reversible dynamics. The spectrum of this matrix is, however, not easy to analyze in general.

Once again, *K*(*t*) and *W*_2_(*t*) are easy to compute thanks to the fact that **C**(*t*) commutes with **∑**, yielding

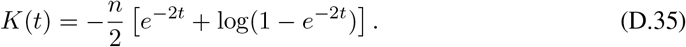

and

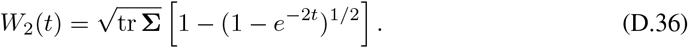

Comparing the *W*_2_ distance for naïve Langevin sampling (D.23) with (D.36), we can see that sampling in the space of natural parameters eliminates the sensitivity of convergence speed to large eigenvalues of the covariance matrix. Moreover, comparing the stationary cross-covariance at timelag *𝒯*≡ *t*− *t*^*′*^ for naïve Langevin sampling, 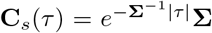, to that for sampling in the natural space, **C**_*s*_(*𝒯*) = *e*^−|*𝒯*|^**∑**, we can see that the same qualitative difference is present. We note that Hennequin et al. [9] focused on the speed of decay in **C**_*s*_(*𝒯*). Therefore, in this simple setting, there is a clear intuitive picture of why natural gradients enable fast sampling.

## E Numerical methods and supplementary figures

In this appendix, we describe our numerical methods and include supplementary figures. All simulations were run in Matlab 9.10 (R2021a) or 9.12 (R2022a) (The MathWorks, Natick, MA, USA) on desktop workstations (CPU: Intel i9-9900K or Xeon W-2145, 64GB RAM). They were not computationally intensive, and required less than 24 hours of compute time in total. The code used to generate all figures is available from GitHub: https://github.com/Pehlevan-Group/FastSpikingSampler.

For the sweeps in *ρ* we tested 100 values of *ρ* ∈ [0; 0.99]. For the dimension sweeps we varied *n*_*p*_ ∈ [2, 4, 8, 16, 32, 64] and 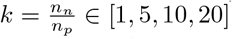 is the number of neurons per parameters. Here, we showed results for *k* = 10. As expected, sampling failed in the case of *k* = 1 as the sign constraint introduced by spiking restricts the network to efficiently sample only one half on the values for each parameter but results were qualitatively similar for *k* = *n*_*n*_*/n*_*p*_ ∈ [5, 10, 20].

In Figures 2, E.2, E.3, 4, and E.4, convergence statistics are computed based on distributions of samples over time, that is, the values visited by the sampler over the course of a single trial. Note that this is different than many machine learning studies, which instead consider distributions across realizations at a single timepoint. However, for an organism, probabilistic inference must be performed within a single trial. Because estimation of the full 2-Wasserstein distance between high-dimensional distributions is computationally expensive [98], we instead computed the mean across dimensions of the 2-Wasserstein distances between the marginals of the sampling and target distributions.

For Figures 2, E.2 and E.3, we sampled with a discretization timestep of Δ = 10^−4^ s and a membrane time constant of *𝒯*_*m*_ = 20 ms. **Γ** by drawing independent and identically distributed Gaussian element. We uses *𝒯*_*s*_ = 0.01*𝒯*_*m*_ and scaled the elastic net regularization parameters using 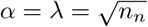.

For Figures 3, 4, and E.4, we simulated a Metropolis-Hastings sampling network with a discretization timestep of Δ = 10^−5^ s and a membrane time constant of *𝒯*_*m*_ = 20 ms. In the naïve case, we generated the readout matrices as **Γ** = [− **M, M**] for random matrices **M** with independent and identically distributed Gaussian elements. To perform sampling in approximately the natural space, we chose **Γ** = **∑**^1*/*2^[−**M, M**] for **M** a random matrix with i.i.d. Gaussian elements. For the dimension sweeps in 4c and E.4b, we scale the variance of the elements of **M** to be 1*/n*_*p*_. We probe the sensitivity of the sampler to the variance of the random matrix in Figure B.1, showing that its performance is robust within some range of variances. The distributions in Figure 3b were generated using 1000 realizations of the randomness in the proposal and accept/reject steps for a single realization of the random matrix **M**. Statistics in Figures 4 and E.4 were computed across 100 realizations of the random matrix **M** and of the randomness in the proposal and accept/reject steps.

**Figure E.1.**
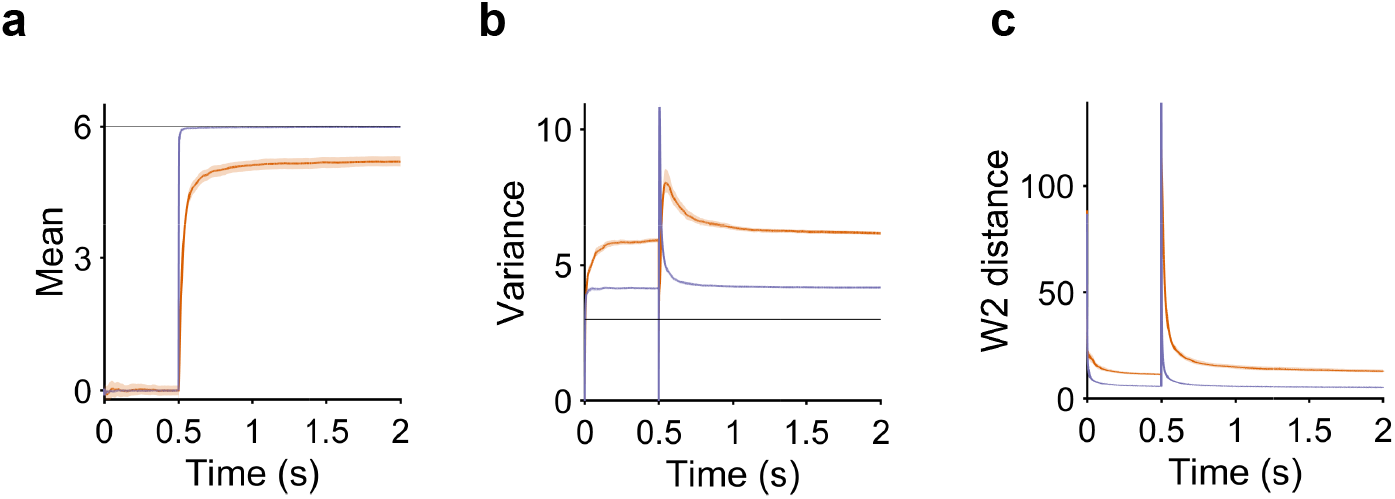
Time course of the inference statistics in EBNs implementing Langevin sampling using naïve and natural geometry. There is initially no input (target mean = 0). At time *t* = 0.5*s*, the stimulus appears (target mean =6). The network parameters are *ρ* = 0.8, *n*_*p*_ = 20 and *n*_*n*_ = 200. This corresponds to network parameters for which the difference between the naïve and natural geometry starts to be sizeable (See Figures 2.c and E.3.a). **a**. The inferred mean converges faster for the natural geometry and there is a steady state error in the naïve implementation. **b**. The transient in the value of the estimated variance at stimulus onset (*t* = 0.5*s*) and the steady state error are larger and the transient relaxes to baseline more slowly in the naïve implementation than with the natural geometry. **c**. The transient in the *W*_2_ distance to the target distribution at stimulus onset (*t* = 0.5*s*) and the steady state *W*_2_ distance are larger and the transient relaxes to baseline more slowly in the naïve implementation than with the natural geometry. Shaded error patches show 95% confidence intervals computed via bootstrapping over 100 realizations in all panels

**Figure E.2.**
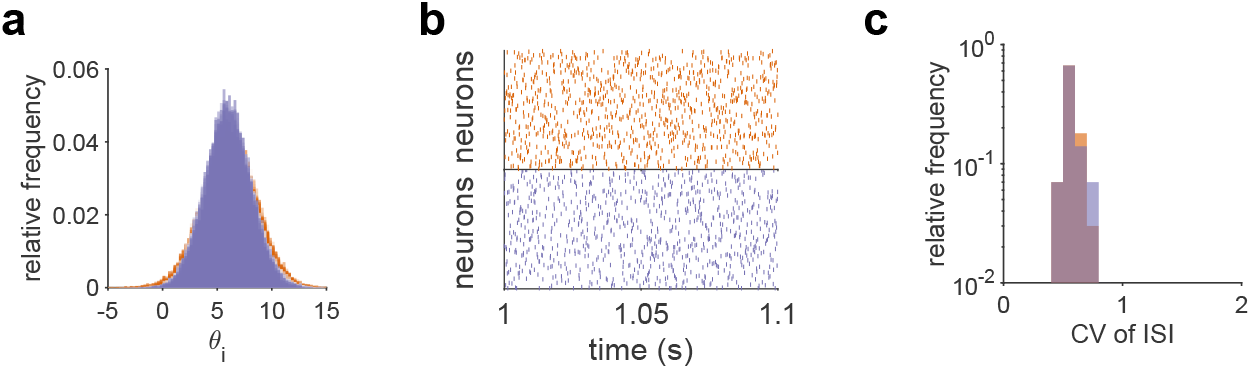
Additional statistics of spike trains in EBNs implementing Langevin sampling using naïve and natural geometry. **a**. Marginal distribution of *θ*_*i*_ after stimulus onset. **b**. Example raster plots over a 100 ms window for naïve and natural geometry. **c**. The distribution of coefficients of variation of ISIs across neurons.

**Figure E.3.**
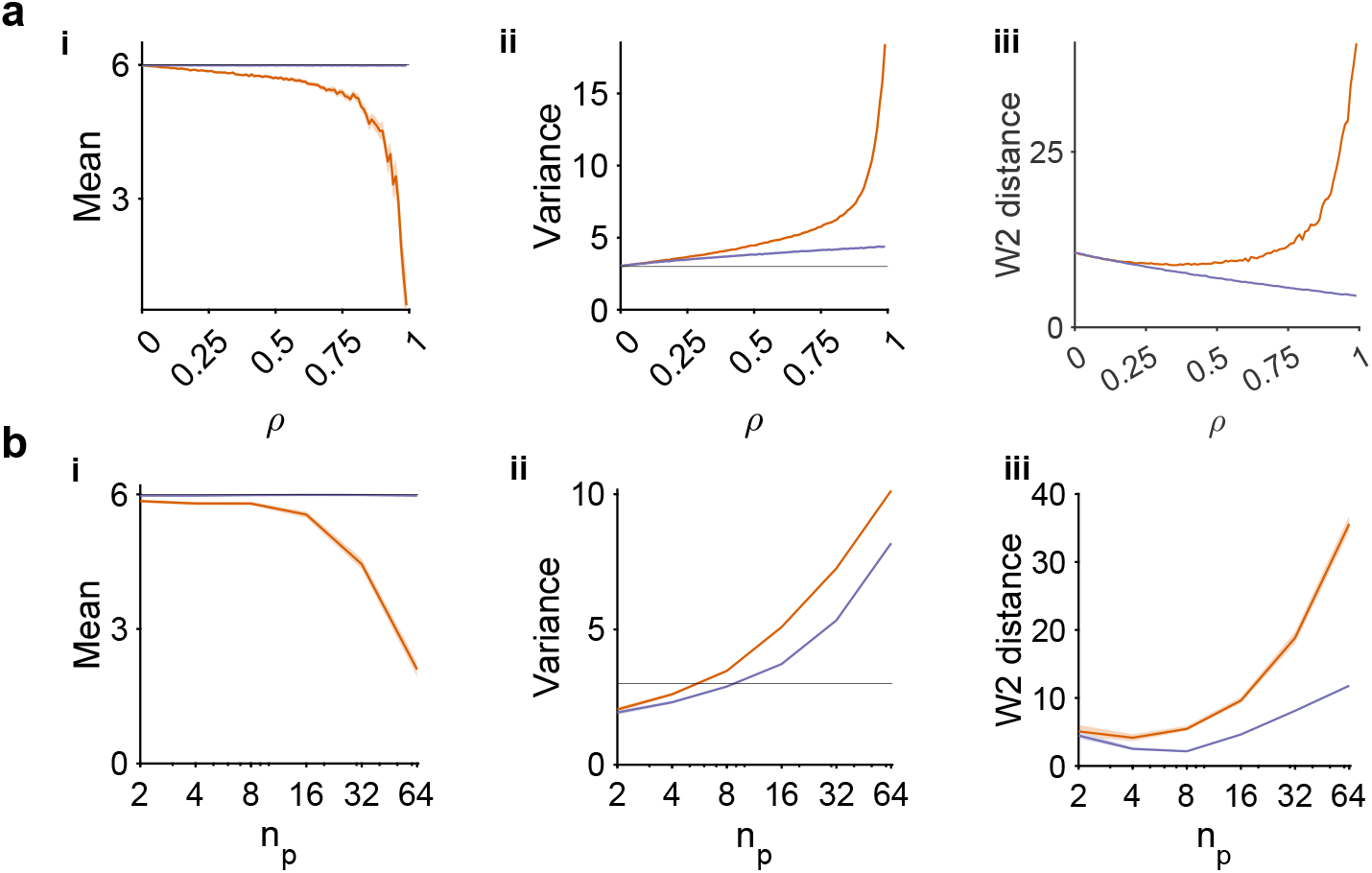
Inference statistics for sampling in efficient balanced networks at steady state (using full 1.5s after stimulus onset). The setup is as in Figure 2. **a**. Comparison of performance in the 50 ms following stimulus onset (after *t* = 0.5 s) between naïve and natural geometry for varying *ρ* **a.i**. The estimate of the mean collapses towards zero with increasing *ρ* for naïve geometry, **a.ii**. The estimated variance increases catastrophically with *ρ* for naïve geometry, but only mildly for natural geometry. **a.iii**. Inference accuracy as measured by the mean marginal 2-Wasserstein distance across dimensions decreases for naïve. **b**. Similar analysis when varying *n*_*p*_ for the **d.i**. Mean, **d.ii**. Variance and **b.iii**. mean 2-Wasserstein distance. Shaded error patches show 95% confidence intervals computed via bootstrapping over 100 realizations in all panels; see Appendix E for further details. See Appendix E for detailed numerical methods.

**Figure E.4.**
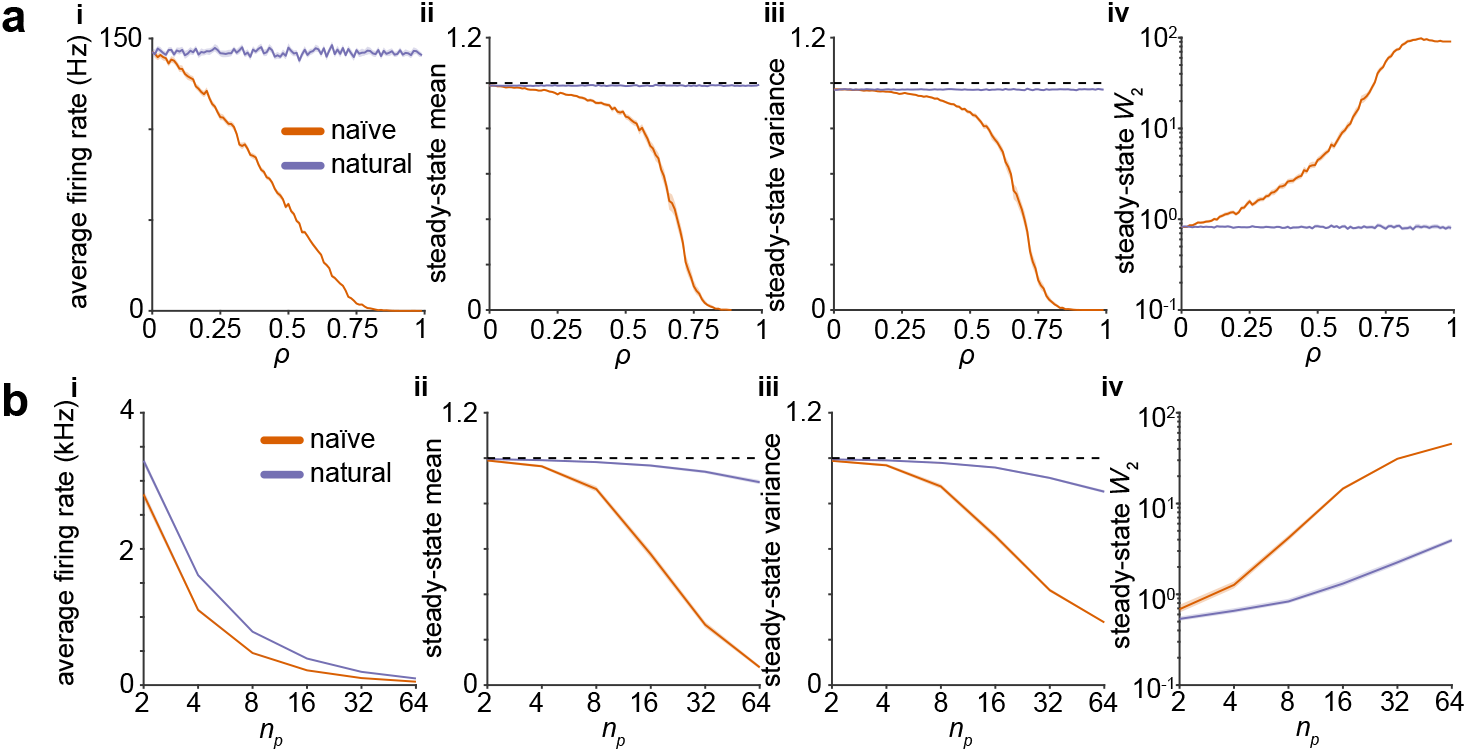
Steady-state statistics for the Metropolis-Hastings sampler of Figure 3. **a**. Sampling from strongly-equicorrelated Gaussians in *n*_*p*_ = 10 dimensions using a network of *n*_*n*_ = 100 neurons requires careful choice of geometry. The stimulus setup is as in Figure 3. **a.i**. At strong correlations *ρ*, spiking is suppressed in networks with naïve geometry, but not when the natural geometry is used. This panel is identical to Figure 3b.i. **a.ii**. Dimension-averaged estimate of the mean signal over the entire stimulus interval. **a.iii**. As in **ii**, but for the variance. **a.iv**. As in **ii**, but for the 2-Wasserstein distance. **b**. Sampling in high dimensions requires careful choice of geometry. **b.i**. At moderately strong correlations *ρ* = 0.75 and high dimensions, spiking is suppressed in networks with naïve geometry, but not when the natural geometry is used. Here, we use 10 neurons per parameter, i.e., *n*_*n*_ = 10*n*_*p*_. **b.ii**. Dimension-averaged estimate of the mean signal in the 50 milliseconds after stimulus onset. This panel is identical to Figure 3c.i. **b.iii**. As in **ii**, but for the variance. **b.iv**. As in **ii**, but for the 2-Wasserstein distance. Shaded error patches show 95% confidence intervals computed via bootstrapping over 100 realizations in all panels; see Appendix E for further details.

## Footnotes

3 In Appendix B.3, we show how the elastic net prior on firing rates used by Boerlin et al. [45] can be incorporated into this model. As it only modifies the definitions of the recurrent weights and spiking threshold, this extension does not add new conceptual difficulties, hence we do not discuss it further in the main text. Additionally, we provide a pedagogical introduction to the dynamics of the EBN in Appendix C.

4 In Appendix C, we give a concrete interpretation of this task in terms of the Gaussian linear models used in previous work by Hennequin et al. [9] and Savin and Denève [20].

5 Here, we chose to include the regularizing term *α* as a fixed offset in the voltage potential but it can equivalently be included as an offset in the spiking threshold *T*_*i*_, as discussed in Appendix B.3.

## References

[1] David C Knill and Alexandre Pouget. The Bayesian brain: the role of uncertainty in neural coding and computation. Trends in Neurosciences, 27(12):712–719, 2004. doi: 10.1016/j.tins.2004.10.007.

[2] Konrad P Körding and Daniel M Wolpert. Bayesian integration in sensorimotor learning. Nature, 427(6971):244–247, 2004. doi: 10.1038/nature02169.

[3] József Fiser, Pietro Berkes, Gergoő Orbán, and Máté Lengyel. Statistically optimal perception and learning: from behavior to neural representations. Trends in Cognitive Sciences, 14(3): 119–130, 2010. doi: 10.1016/j.tics.2010.01.003.

[4] Torben Ott, Paul Masset, and Adam Kepecs. The neurobiology of confidence: From beliefs to neurons. In Cold Spring Harbor Symposia on Quantitative Biology, volume 83, pages 9–16. Cold Spring Harbor Laboratory Press, 2018. doi: 10.1101/sqb.2018.83.038794.

[5] Paul Masset, Torben Ott, Armin Lak, Junya Hirokawa, and Adam Kepecs. Behavior-and modality-general representation of confidence in orbitofrontal cortex. Cell, 182(1):112–126, 2020. doi: 10.1016/j.cell.2020.05.022.

[6] Ruben S Van Bergen, Wei Ji Ma, Michael S Pratte, and Janneke FM Jehee. Sensory uncertainty decoded from visual cortex predicts behavior. Nature Neuroscience, 18(12):1728–1730, 2015. doi: 10.1038/nn.4150.

[7] Dylan Festa, Amir Aschner, Aida Davila, Adam Kohn, and Ruben Coen-Cagli. Neuronal vari-ability reflects probabilistic inference tuned to natural image statistics. Nature Communications, 12(1):1–11, 2021. doi: 10.1038/s41467-021-23838-x.

[8] Alexandre Pouget, Jan Drugowitsch, and Adam Kepecs. Confidence and certainty: distinct probabilistic quantities for different goals. Nature Neuroscience, 19(3):366–374, 2016. doi: 10.1038/nn.4240.

[9] Guillaume Hennequin, Laurence Aitchison, and Mate Lengyel. Fast sampling-based inference in balanced neuronal networks. In Z. Ghahramani, M. Welling, C. Cortes, N. Lawrence, and K.Q. Weinberger, editors, Advances in Neural Information Processing Systems, volume 27, pages 2240–2248. Curran Associates, Inc., 2014. URL https://papers.nips.cc/paper/2014/hash/a7d8ae4569120b5bec12e7b6e9648b86-Abstract.html.

[10] Alexandre Pouget, Jeffrey M Beck, Wei Ji Ma, and Peter E Latham. Probabilistic brains: knowns and unknowns. Nature Neuroscience, 16(9):1170–1178, 2013. doi: 10.1038/nn.3495.

[11] Ádám Koblinger, József Fiser, and Máté Lengyel. Representations of uncertainty: where art thou? Current Opinion in Behavioral Sciences, 38:150–162, 2021. doi: 10.1016/j.cobeha.2021.03.009.

[12] Hansem Sohn and Devika Narain. Neural implementations of Bayesian inference. Current Opinion in Neurobiology, 70:121–129, 2021. doi: 10.1016/j.conb.2021.09.008.

[13] Wei Ji Ma, Jeffrey M Beck, Peter E Latham, and Alexandre Pouget. Bayesian inference with probabilistic population codes. Nature Neuroscience, 9(11):1432–1438, 2006. doi: 10.1038/nn1790.

[14] Maneesh Sahani and Peter Dayan. Doubly distributional population codes: simultaneous representation of uncertainty and multiplicity. Neural Computation, 15(10):2255–2279, 2003. doi: 10.1162/089976603322362356.

[15] Eszter Vértes and Maneesh Sahani. Flexible and accurate inference and learning for deep generative models. In S. Bengio, H. Wallach, H. Larochelle, K. Grauman, N. Cesa-Bianchi, and R. Garnett, editors, Advances in Neural Information Processing Systems, volume 31. Curran Associates, Inc., 2018. URL https://proceedings.neurips.cc/paper/2018/hash/955cb567b6e38f4c6b3f28cc857fc38c-Abstract.html.

[16] Patrik Hoyer and Aapo Hyvärinen. Interpreting neural response variability as Monte Carlo sampling of the posterior. In S. Becker, S. Thrun, and K. Ober-mayer, editors, Advances in Neural Information Processing Systems, volume 15. MIT Press, 2002. URL https://proceedings.neurips.cc/paper/2002/hash/a486cd07e4ac3d270571622f4f316ec5-Abstract.html.

[17] Lars Buesing, Johannes Bill, Bernhard Nessler, and Wolfgang Maass. Neural dynamics as sampling: a model for stochastic computation in recurrent networks of spiking neurons. PLoS Computational Biology, 7(11):e1002211, 2011. doi: 10.1371/journal.pcbi.1002211.

[18] Dejan Pecevski, Lars Buesing, and Wolfgang Maass. Probabilistic inference in general graphical models through sampling in stochastic networks of spiking neurons. PLOS Computational Biology, 7(12):1–25, 12 2011. doi: 10.1371/journal.pcbi.1002294. URL https://doi.org/10.1371/journal.pcbi.1002294.

[19] Agnieszka Grabska-Barwinska, Jeff Beck, Alexandre Pouget, and Peter Latham. Demixing odors - fast inference in olfaction. In C.J. Burges, L. Bottou, M. Welling, Z. Ghahramani, and K.Q. Weinberger, editors, Advances in Neural Information Processing Systems, volume 26, pages 1968–1976. Curran Associates, Inc., 2013. URL https://proceedings.neurips.cc/paper/2013/hash/2bcab9d935d219641434683dd9d18a03-Abstract.html.

[20] Cristina Savin and Sophie Denève. Spatio-temporal representations of uncertainty in spiking neural networks. In Z. Ghahramani, M. Welling, C. Cortes, N. Lawrence, and K.Q. Weinberger, editors, Advances in Neural Information Processing Systems, volume 27, pages 2024–2032. Curran Associates, Inc., 2014. URL https://proceedings.neurips.cc/paper/2014/hash/4e2545f819e67f0615003dd7e04a6087-Abstract.html.

[21] Laurence Aitchison and Máté Lengyel. The Hamiltonian brain: Efficient probabilistic inference with excitatory-inhibitory neural circuit dynamics. PLoS Computational Biology, 12(12): e1005186, 2016. doi: 10.1371/journal.pcbi.1005186.

[22] Wen-Hao Zhang, Tai Sing Lee, Brent Doiron, and Si Wu. Distributed sampling-based Bayesian inference in coupled neural circuits. bioRxiv, 2020. doi: 10.1101/2020.07.20.212126.

[23] Rodrigo Echeveste, Laurence Aitchison, Guillaume Hennequin, and Máté Lengyel. Cortical-like dynamics in recurrent circuits optimized for sampling-based probabilistic inference. Nature Neuroscience, 23(9):1138–1149, 2020. doi: 10.1038/s41593-020-0671-1.

[24] Wenhao Zhang, Si Wu, Kresimir Josic, and Brent Doiron. Sampling-based Bayesian inference in recurrent circuits of stochastic spiking neurons. bioRxiv, 2022. doi: 10.1101/2022.01.26.477877.

[25] Yang Qi and Pulin Gong. Fractional neural sampling as a theory of spatiotemporal probabilistic computations in neural circuits. Nature Communications, 13(1):4572, Aug 2022. ISSN 2041-1723. doi: 10.1038/s41467-022-32279-z.

[26] Jeffrey M. Beck, Peter E. Latham, and Alexandre Pouget. Marginalization in neural circuits with divisive normalization. Journal of Neuroscience, 31(43):15310–15319, 2011. ISSN 0270-6474. doi: 10.1523/JNEUROSCI.1706-11.2011. URL https://www.jneurosci.org/content/31/43/15310.

[27] Radford M Neal. Probabilistic inference using Markov chain Monte Carlo methods. Department of Computer Science, University of Toronto Toronto, ON, Canada, 1993. URL https://www.cs.toronto.edu/~radford/review.abstract.html.

[28] Christian P Robert, Víctor Elvira, Nick Tawn, and Changye Wu. Accelerating MCMC algo-rithms. Wiley Interdisciplinary Reviews: Computational Statistics, 10(5):e1435, 2018. doi: 10.1002/wics.1435.

[29] Crispin W Gardiner. Handbook of stochastic methods, volume 3. Springer Berlin, 1985.

[30] Max Welling and Yee W Teh. Bayesian learning via stochastic gradient Langevin dynamics. In Proceedings of the 28th international conference on machine learning (ICML-11), pages 681–688, 2011. URL https://icml.cc/2011/papers/398_icmlpaper.pdf.

[31] Radford M Neal. MCMC using Hamiltonian dynamics. In Handbook of Markov Chain Monte Carlo, pages 139–188. Chapman and Hall/CRC, 2011. doi: 10.1201/b10905.

[32] Alain Durmus and Eric Moulines. Nonasymptotic convergence analysis for the unadjusted Langevin algorithm. The Annals of Applied Probability, 27(3):1551–1587, 2017. doi: 10.1214/16-AAP1238.

[33] Santosh Vempala and Andre Wibisono. Rapid convergence of the unadjusted Langevin algo-rithm: Isoperimetry suffices. In H. Wallach, H. Larochelle, A. Beygelzimer, F. d’Alché-Buc, E. Fox, and R. Garnett, editors, Advances in Neural Information Processing Systems, volume 32. Curran Associates, Inc., 2019. URL https://proceedings.neurips.cc/paper/2019/hash/65a99bb7a3115fdede20da98b08a370f-Abstract.html.

[34] Sam Patterson and Yee Whye Teh. Stochastic gradient Riemannian Langevin dynamics on the probability simplex. In C.J. Burges, L. Bottou, M. Welling, Z. Ghahramani, and K.Q. Weinberger, editors, Advances in Neural Information Processing Systems, volume 26. Curran Associates, Inc., 2013. URL https://proceedings.neurips.cc/paper/2013/hash/309928d4b100a5d75adff48a9bfc1ddb-Abstract.html.

[35] Pavel Izmailov, Sharad Vikram, Matthew D Hoffman, and Andrew Gordon Gordon Wilson. What are Bayesian neural network posteriors really like? In International Conference on Ma-chine Learning, pages 4629–4640. PMLR, 2021. URL https://proceedings.mlr.press/v139/izmailov21a.html.

[36] Florian Wenzel, Kevin Roth, Bastiaan Veeling, Jakub Swiatkowski, Linh Tran, Stephan Mandt, Jasper Snoek, Tim Salimans, Rodolphe Jenatton, and Sebastian Nowozin. How good is the Bayes posterior in deep neural networks really? In Hal Daumé III and Aarti Singh, editors, Proceedings of the 37th International Conference on Machine Learning, volume 119 of Proceedings of Machine Learning Research, pages 10248–10259. PMLR, 13–18 Jul 2020. URL https://proceedings.mlr.press/v119/wenzel20a.html.

[37] Nikolaus Kriegeskorte and Xue-Xin Wei. Neural tuning and representational geometry. Nature Reviews Neuroscience, 09 2021. ISSN 1471-0048. doi: 10.1038/s41583-021-00502-3.

[38] SueYeon Chung and LF Abbott. Neural population geometry: An approach for understanding biological and artificial neural networks. Current Opinion in Neurobiology, 70:137–144, 2021. doi: 10.1016/j.conb.2021.10.010.

[39] Shun-Ichi Amari. Natural gradient works efficiently in learning. Neural Computation, 10(2): 251–276, 1998. doi: 10.1162/089976698300017746.

[40] Shun-Ichi Amari and Scott C Douglas. Why natural gradient? In Proceedings of the 1998 IEEE International Conference on Acoustics, Speech and Signal Processing, ICASSP’98 (Cat. No. 98CH36181), volume 2, pages 1213–1216. IEEE, 1998. doi: 10.1109/ICASSP.1998.675489.

[41] Shun-Ichi Amari. Information geometry and its applications, volume 194. Springer, 2016. doi: 10.1007/978-4-431-55978-8.

[42] Mark Girolami and Ben Calderhead. Riemann manifold Langevin and Hamiltonian Monte Carlo methods. Journal of the Royal Statistical Society: Series B (Statistical Methodology), 73 (2):123–214, 2011. doi: 10.1111/j.1467-9868.2010.00765.x.

[43] Yi-An Ma, Tianqi Chen, and Emily Fox. A complete recipe for stochastic gradient MCMC. In C. Cortes, N. Lawrence, D. Lee, M. Sugiyama, and R. Garnett, editors, Advances in Neural Infor-mation Processing Systems, volume 28. Curran Associates, Inc., 2015. URL https://papers.nips.cc/paper/2015/hash/9a4400501febb2a95e79248486a5f6d3-Abstract.html.

[44] Christopher Nemeth and Paul Fearnhead. Stochastic gradient Markov chain Monte Carlo. Journal of the American Statistical Association, 116(533):433–450, 2021. doi: 10.1080/01621459.2020.1847120.

[45] Martin Boerlin, Christian K Machens, and Sophie Denève. Predictive coding of dynamical variables in balanced spiking networks. PLoS Computational Biology, 9(11):e1003258, 2013. doi: 10.1371/journal.pcbi.1003258.

[46] Sophie Denève and Christian K Machens. Efficient codes and balanced networks. Nature Neuroscience, 19(3):375–382, 2016. doi: 10.1038/nn.4243.

[47] Sophie Denève, Alireza Alemi, and Ralph Bourdoukan. The brain as an efficient and robust adaptive learner. Neuron, 94(5):969–977, 2017. doi: 10.1016/j.neuron.2017.05.016.

[48] Camille E Rullán Buxó and Jonathan W Pillow. Poisson balanced spiking networks. PLoS Computational Biology, 16(11):e1008261, 2020. doi: 10.1371/journal.pcbi.1008261.

[49] Nuno Calaim, Florian A Dehmelt, Pedro J Gonçalves, and Christian K Machens. The geometry of robustness in spiking neural networks. eLife, 11:e73276, may 2022. ISSN 2050-084X. doi: 10.7554/eLife.73276. URL https://doi.org/10.7554/eLife.73276.

[50] W. Keith Hastings. Monte Carlo sampling methods using Markov chains and their applications. Biometrika, 57(1):97–109, 1970. doi: 10.1093/biomet/57.1.97.

[51] Radford M. Neal. Improving asymptotic variance of MCMC estimators: Non-reversible chains are better. arXiv, 2004. doi: 10.48550/ARXIV.MATH/0407281. URL https://arxiv.org/abs/math/0407281.

[52] Bernt Øksendal. Stochastic differential equations. Springer, 2003. doi: 10.1007/978-3-642-14394-6.

[53] Michael Betancourt. A conceptual introduction to Hamiltonian Monte Carlo. arXiv preprint arXiv:1701.02434, 2017. doi: 10.48550/arXiv.1701.02434.

[54] Zongchen Chen and Santosh S Vempala. Optimal convergence rate of Hamiltonian Monte Carlo for strongly logconcave distributions. Theory of Computing, 18(1):1–18, 2022. doi: 10.4086/toc.2022.v018a009.

[55] Nan Ding, Youhan Fang, Ryan Babbush, Changyou Chen, Robert D Skeel, and Hartmut Neven. Bayesian sampling using stochastic gradient thermostats. In Z. Ghahramani, M. Welling, C. Cortes, N. Lawrence, and K.Q. Weinberger, edi-tors, Advances in Neural Information Processing Systems, volume 27. Curran As-sociates, Inc., 2014. URL https://proceedings.neurips.cc/paper/2014/hash/21fe5b8ba755eeaece7a450849876228-Abstract.html.

[56] Jianghong Shi, Tianqi Chen, Ruoshi Yuan, Bo Yuan, and Ping Ao. Relation of a new interpreta-tion of stochastic differential equations to Ito process. Journal of Statistical Physics, 148(3): 579–590, 2012. doi: 10.1007/s10955-012-0532-8.

[57] Razvan Pascanu and Yoshua Bengio. Revisiting natural gradient for deep networks. arXiv preprint arXiv:1301.3584, 2013. doi: 10.48550/arXiv.1301.3584.

[58] James Martens. New insights and perspectives on the natural gradient method. Journal of Machine Learning Research, 21:1–76, 2020. URL https://jmlr.org/papers/v21/17-678.html.

[59] Jascha Sohl-Dickstein, Mayur Mudigonda, and Michael DeWeese. Hamiltonian Monte Carlo without detailed balance. In Eric P. Xing and Tony Jebara, editors, Proceedings of the 31st International Conference on Machine Learning, volume 32 of Proceedings of Ma-chine Learning Research, pages 719–726, Bejing, China, 22–24 Jun 2014. PMLR. URL https://proceedings.mlr.press/v32/sohl-dickstein14.html.

[60] Thomas Rippl, Axel Munk, and Anja Sturm. Limit laws of the empirical Wasserstein distance: Gaussian distributions. Journal of Multivariate Analysis, 151:90–109, 2016. ISSN 0047-259X. doi: https://doi.org/10.1016/j.jmva.2016.06.005. URL https://www.sciencedirect.com/science/article/pii/S0047259X16300446.

[61] William R Softky and Christof Koch. The highly irregular firing of cortical cells is inconsistent with temporal integration of random epsps. Journal of Neuroscience, 13(1):334–350, 1993. doi: 10.1523/JNEUROSCI.13-01-00334.1993.

[62] Carl van Vreeswijk and Haim Sompolinsky. Chaotic balanced state in a model of cortical circuits. Neural Computation, 10(6):1321–1371, 1998. doi: 10.1162/089976698300017214.

[63] Christopher De Sa, Chris Re, and Kunle Olukotun. Ensuring rapid mixing and low bias for asynchronous Gibbs sampling. In International Conference on Machine Learning, pages 1567–1576. PMLR, 2016. URL https://proceedings.mlr.press/v48/sa16.html.

[64] Alexander Terenin, Daniel Simpson, and David Draper. Asynchronous Gibbs sampling. In International Conference on Artificial Intelligence and Statistics, pages 144–154. PMLR, 2020. URL https://proceedings.mlr.press/v108/terenin20a.html.

[65] Nicolas Brunel. Dynamics of sparsely connected networks of excitatory and inhibitory spiking neurons. Journal of Computational Neuroscience, 8(3):183–208, 2000. doi: 10.1023/a:1008925309027.

[66] Alfonso Renart, Nicolas Brunel, and Xiao-Jing Wang. Mean-field theory of irregularly spiking neuronal populations and working memory in recurrent cortical networks. Computational Neuroscience: A comprehensive approach, pages 431–490, 2004. doi: 10.1201/9780203494462-22.

[67] Alexander Lerchner, Cristina Ursta, John Hertz, Mandana Ahmadi, Pauline Ruffiot, and Søren Enemark. Response variability in balanced cortical networks. Neural Computation, 18(3): 634–659, 2006. doi: 10.1162/089976606775623261.

[68] Gareth O Roberts and Richard L Tweedie. Exponential convergence of Langevin distributions and their discrete approximations. Bernoulli, pages 341–363, 1996. doi: 10.2307/3318418.

[69] Ruqi Zhang, A Feder Cooper, and Christopher De Sa. AMAGOLD: Amortized Metropolis adjustment for efficient stochastic gradient MCMC. In International Conference on Artificial Intelligence and Statistics, pages 2142–2152. PMLR, 2020. URL https://proceedings.mlr.press/v108/zhang20e.html.

[70] Michele Nardin, James W Phillips, William F Podlaski, and Sander W Keemink. Nonlinear computations in spiking neural networks through multiplicative synapses. Peer Community Journal, 1, 2021. doi: 10.24072/pcjournal.69.

[71] Mengchen Zhu and Christopher J Rozell. Modeling inhibitory interneurons in efficient sensory coding models. PLoS Computational Biology, 11(7):e1004353, 2015. doi: 10.1371/jour-nal.pcbi.1004353.

[72] Arish Alreja, Ilya Nemenman, and Christopher J Rozell. Constrained brain volume in an efficient coding model explains the fraction of excitatory and inhibitory neurons in sensory cortices. PLOS Computational Biology, 18(1):e1009642, 2022. doi: 10.1371/journal.pcbi.1009642.

[73] Jose Del Castillo and Bernard Katz. Quantal components of the end-plate potential. The Journal of Physiology, 124(3):560, 1954. doi: 10.1113/jphysiol.1954.sp005129.

[74] Rob R de Ruyter van Steveninck, Geoffrey D Lewen, Steven P Strong, Roland Koberle, and William Bialek. Reproducibility and variability in neural spike trains. Science, 275(5307): 1805–1808, 1997. doi: 10.1126/science.275.5307.1805.

[75] Gianluigi Mongillo, Omri Barak, and Misha Tsodyks. Synaptic theory of working memory. Science, 319(5869):1543–1546, 2008. doi: 10.1126/science.1150769.

[76] Dmitri A Rusakov, Leonid P Savtchenko, and Peter E Latham. Noisy synaptic conductance: Bug or a feature? Trends in Neurosciences, 43(6):363–372, 2020. doi: 10.1016/j.tins.2020.03.009.

[77] Elena Kreutzer, Walter Senn, and Mihai A Petrovici. Natural-gradient learning for spiking neurons. eLife, 11:e66526. 05 2022. ISSN 2050-084X. doi: 10.7554/eLife.66526.

[78] Kwabena Boahen. A neuromorph’s prospectus. Computing in Science & Engineering, 19(2): 14–28, 2017. doi: 10.1109/MCSE.2017.33.

[79] Mike Davies, Narayan Srinivasa, Tsung-Han Lin, Gautham Chinya, Yongqiang Cao, Sri Harsha Choday, Georgios Dimou, Prasad Joshi, Nabil Imam, Shweta Jain, et al. Loihi: A neuro-morphic manycore processor with on-chip learning. IEEE Micro, 38(01):82–99, 2018. doi: 10.1109/MM.2018.112130359.

[80] Kaushik Roy, Akhilesh Jaiswal, and Priyadarshini Panda. Towards spike-based machine intelligence with neuromorphic computing. Nature, 575(7784):607–617, 2019. doi: https://doi.org/10.1038/s41586-019-1677-2.

[81] Benjamin Cramer, Sebastian Billaudelle, Simeon Kanya, Aron Leibfried, Andreas Grübl, Vitali Karasenko, Christian Pehle, Korbinian Schreiber, Yannik Stradmann, Johannes Weis, et al. Surrogate gradients for analog neuromorphic computing. Proceedings of the National Academy of Sciences, 119(4), 2022. doi: 10.1073/pnas.2109194119.

[82] David Kappel, Stefan Habenschuss, Robert Legenstein, and Wolfgang Maass. Network plas-ticity as Bayesian inference. PLoS Computational Biology, 11(11):e1004485, 2015. doi: 10.1371/journal.pcbi.1004485.

[83] Laurence Aitchison, Jannes Jegminat, Jorge Aurelio Menendez, Jean-Pascal Pfister, Alexandre Pouget, and Peter E Latham. Synaptic plasticity as Bayesian inference. Nature Neuroscience, 24(4):565–571, 2021. doi: 10.1038/s41593-021-00809-5.

[84] Paul Masset, Shanshan Qin, and Jacob A Zavatone-Veth. Drifting neuronal representations: Bug or feature? Biological Cybernetics, pages 1–14, 2022. doi: 10.1007/s00422-021-00916-3.

[85] Wenbo Gong, Yingzhen Li, and José Miguel Hernández-Lobato. Meta-learning for stochastic gradient MCMC. In 7th International Conference on Learning Representations, ICLR 2019, 2019. doi: 10.48550/arXiv.1806.04522.

[86] Jacob A. Zavatone-Veth, Abdulkadir Canatar, Benjamin S. Ruben, and Cengiz Pehlevan. Asymptotics of representation learning in finite Bayesian neural networks. In M. Ran-zato, A. Beygelzimer, Y. Dauphin, P.S. Liang, and J. Wortman Vaughan, editors, Ad-vances in Neural Information Processing Systems, volume 34, pages 24765–24777. Curran Associates, Inc., 2021. URL https://proceedings.neurips.cc/paper/2021/hash/cf9dc5e4e194fc21f397b4cac9cc3ae9-Abstract.html.

[87] Ralph Bourdoukan, David Barrett, Sophie Deneve, and Christian K Machens. Learning op-timal spike-based representations. In F. Pereira, C.J. Burges, L. Bottou, and K.Q. Wein-berger, editors, Advances in Neural Information Processing Systems, volume 25. Curran Associates, Inc., 2012. URL https://proceedings.neurips.cc/paper/2012/hash/3a15c7d0bbe60300a39f76f8a5ba6896-Abstract.html.

[88] Alireza Alemi, Christian Machens, Sophie Denève, and Jean-Jacques Slotine. Learning nonlinear dynamics in efficient, balanced spiking networks using local plasticity rules. In Proceedings of the AAAI Conference on Artificial Intelligence, volume 32, 2018. doi: 10.1609/aaai.v32i1.11320.

[89] Wieland Brendel, Ralph Bourdoukan, Pietro Vertechi, Christian K Machens, and Sophie Denéve. Learning to represent signals spike by spike. PLoS Computational Biology, 16(3):e1007692, 2020. doi: 10.1371/journal.pcbi.1007692.

[90] Dongsung Huh and Terrence J Sejnowski. Gradient descent for spiking neural networks. In S. Bengio, H. Wallach, H. Larochelle, K. Grauman, N. Cesa-Bianchi, and R. Gar-nett, editors, Advances in Neural Information Processing Systems, volume 31. Curran Associates, Inc., 2018. URL https://proceedings.neurips.cc/paper/2018/hash/185e65bc40581880c4f2c82958de8cfe-Abstract.html.

[91] Emre O Neftci, Hesham Mostafa, and Friedemann Zenke. Surrogate gradient learning in spiking neural networks: Bringing the power of gradient-based optimization to spiking neural networks. IEEE Signal Processing Magazine, 36(6):51–63, 2019. doi: 10.1109/MSP.2019.2931595.

[92] Cengiz Pehlevan. A spiking neural network with local learning rules derived from non-negative similarity matching. In ICASSP 2019 - 2019 IEEE International Conference on Acoustics, Speech and Signal Processing (ICASSP), pages 7958–7962, 2019. doi: 10.1109/ICASSP.2019.8682290.

[93] Nicolas Perez-Nieves, Vincent CH Leung, Pier Luigi Dragotti, and Dan FM Goodman. Neural heterogeneity promotes robust learning. Nature Communications, 12(1):1–9, 2021. doi: 10.1038/s41467-021-26022-3.

[94] Roger A Horn and Charles R Johnson. Matrix Analysis. Cambridge University Press, 2012. doi: https://doi.org/10.1017/CBO9780511810817.

[95] James Martens and Roger Grosse. Optimizing neural networks with Kronecker-factored approximate curvature. In International Conference on Machine Learning, pages 2408–2417. PMLR, 2015. doi: 10.48550/arXiv.1503.05671.

[96] Chii-Ruey Hwang, Shu-Yin Hwang-Ma, and Shuenn-Jyi Sheu. Accelerating diffusions. The Annals of Applied Probability, 15(2):1433–1444, 2005. doi: 10.1214/105051605000000025.

[97] Claude Godrèche and Jean-Marc Luck. Characterising the nonequilibrium stationary states of Ornstein–Uhlenbeck processes. Journal of Physics A: Mathematical and Theoretical, 52(3): 035002, 2018. doi: 10.1088/1751-8121/aaf190.

[98] Lénaïc Chizat, Pierre Roussillon, Flavien Léger, François-Xavier Vialard, and Gabriel Peyré. Faster Wasserstein distance estimation with the Sinkhorn divergence. In H. Larochelle, M. Ranzato, R. Hadsell, M.F. Balcan, and H. Lin, editors, Advances in Neural Information Processing Systems, volume 33, pages 2257–2269. Curran As-sociates, Inc., 2020. URL https://proceedings.neurips.cc/paper/2020/hash/17f98ddf040204eda0af36a108cbdea4-Abstract.html.

